# Repeat Systemic Delivery of Cross-Neutralization Resistant Synthetic Vesiculoviruses Immunomodulates the Tumor Microenvironment

**DOI:** 10.1101/2025.08.01.668184

**Authors:** Natalie M. Elliott, Benjamin L. Kendall, Souyma Dutta, Daipayan Sarkar, John W. Vant, Jill Thompson, Elizabeth Raupach, Christopher Shin, Khandoker Usran Ferdous, Aleksandra Cios, Kenneth Buetow, Chelsae Watters, Sangkon Oh, Autumn J. Schulze, Alexander T. Baker, Bolni M. Nagalo, Richard Vile, Mitesh J. Borad

**Affiliations:** Department of Molecular Medicine, Mayo Clinic; Rochester, United States; Division of Hematology and Oncology, Mayo Clinic; Phoenix, United States; Department of Immunology, Mayo Clinic; Rochester, United States; School of Molecular Sciences, Arizona State University; Tempe, United States; Laboratory of Chemical Physics, National Institute of Diabetes and Digestive and Kidney Diseases, National Institutes of Health; Bethesda, United States; Computational Sciences and Engineering Division, Oak Ridge National Laboratory; Oak Ridge, United States; Department of Pathology, University of Arkansas for Medical Sciences; Little Rock, United States; Department of Pharmacology and Physiology, University of Maryland School of Medicine; Baltimore, United States; Marlene and Stewart Greenbaum NCI Comprehensive Cancer Cetner, University of Maryland School of Medicine; Baltimore, United States; Computational Sciences and Informatics Program for Complex Adaptive Systems, Arizona State University; Tempe, United States; Joan Reece Professor of Immuno-oncology, Comprehensive Cancer Centre, School of Cancer & Pharmaceutical Sciences, School of Immunology and Microbial Sciences, King’s College London; London, United Kingdom; Mayo Clinic Comprehensive Cancer Center; Phoenix, United States; Center for Individualized Medicine, Mayo Clinic; Rochester, United States

## Abstract

The clinical efficacy of systemic oncolytic virotherapy (OV) is constrained by the rapid development of neutralizing antibodies (nAbs), which prevent repeat systemic administration, a critical barrier to sustained anti-tumor immunity. Vesiculoviruses offer potent oncolytic and immunogenic potential. However, leveraging their serological diversity for repeat dosing remains unexplored. We generated a library of chimeric vesiculovirus vectors incorporating glycoproteins from less well characterized vesiculovirus species. We evaluated vector replication, infectivity, interferon (IFN) responses, and oncolysis *in vitro*, alongside assessments of neutralization resistance using patient sera, monoclonal antibodies, and *in silico* structural modeling. *In vivo* studies assessed tumor delivery, immune activation, and therapeutic efficacy following intravenous administration. The vesiculovirus library exhibited broad tumor infectivity, distinct IFN-stimulatory profiles, and variable oncolytic activity. Neutralization assays and computational modeling identified serological distinctness across vectors, driven by hypervariable glycoprotein epitopes, enabling evasion of cross-neutralizing antibodies. Tumor delivery and anti-tumor immunity were preserved despite humoral barriers. Incorporating tumor-associated antigens (TAAs) further amplified anti-tumor responses, even in the context of anti-viral memory. Sequential administration of distinct vesiculovirus vectors induced robust immune activation and improved survival in a B16-OVA-IFNAR^-/-^ model. Our findings establish a glycoprotein-diverse vesiculovirus platform capable of overcoming humoral immunity, enabling repeat intravenous dosing and sustained engagement of the tumor microenvironment. This strategy advances the field of oncolytic virotherapy by addressing a major translational barrier and lays the groundwork for future clinical studies integrating multi-vector, multi-dose immunovirotherapy with immune checkpoint blockade.

**One Sentence Summary:** A glycoprotein-engineered vesiculovirus platform circumvents neutralizing antibodies, enabling repeat intravenous dosing and sustained anti-tumor immunity in preclinical models.

## Introduction

Oncolytic viruses (OVs) are a promising class of immunotherapeutics engineered to selectively lyse tumor cells while activating proinflammatory signals that promote the recruitment and activation of immune cells in the tumor microenvironment (TME) (*1, 2*). Although early efforts in OV development prioritized viral replication and direct tumor cell lysis, it is now well established that durable responses arise primarily through the induction of systemic anti-tumor immunity (*3*). Mechanistically, systemic or local delivery of OVs promotes inflammatory remodeling of the TME, enabling the expansion and reactivation of tumor-specific T-cells that are often suppressed or exhausted in immunologically “cold” tumors (*4*). However, the rapid development of neutralizing antibodies (nAbs) following initial systemic oncolytic virotherapy remains a major clinical barrier, as patients who relapse after initial tumor control are typically unresponsive to subsequent dosing due to antibody-mediated blockade of viral delivery. This immune barrier represents a critical translational gap, driving efforts to develop OV strategies capable of sustaining TME remodeling and achieving durable anti-tumor responses.

This has led to the adoption of front-loaded dosing regimens (“treatment cycles”) in early-phase trials, designed to deliver multiple doses before the full establishment of adaptive immunity and the generation of neutralizing antibodies. These regimens typically involve successive doses administered early in the treatment course to maximize viral exposure within a permissive immunologic window (*2*). Examples include intravenous administration of oncolytic adenovirus (NCT03852511), coxsackievirus (NCT03408587), and reovirus (NCT00503295) on consecutive days within an initial treatment window. These approaches have demonstrated favorable safety profiles, tumor delivery, and modest anti-tumor activity (*5–8*). Nonetheless, each underscores a fundamental challenge: humoral immunity consistently prevents subsequent systemic OV administration beyond the first cycle, limiting durable therapeutic potential (*9*).

Vesiculoviruses, members of the Rhabdoviridae family, are negative sense, single stranded RNA viruses that have been evaluated as oncolytic viral vectors. Clinically advanced platforms include vesicular stomatitis virus (VSV) and Maraba virus (MARAV). While VSV has been pursued clinically as a single injection therapy(*10–12*), Maraba virus encoding a tumor associated antigen (TAA) has been paired with adenoviral vectors as part of a tumor vaccine prime and boost strategy (*13, 14*). The heterologous prime-boost regimen with two different viruses was employed to evade anti-viral humoral immunity induced by the priming dose, allowing delivery of the second dose of virus to re-expose the adaptive immune system to the tumor antigen carried by the viral vectors (*15–17*). In preclinical studies of repeat dosing, intratumoral delivery with successive doses of the same viral vector enhanced therapeutic efficacy by leveraging pre-existing immunity to boost systemic anti-tumor efficacy (*18–20*). Additional studies illustrate that as long as delivery of the virus to the tumor is successful, an anti-tumor effect is observed even in the absence of viral replication, countering concerns around reduced replication in the presence of anti-viral immunity (*21, 22*). Taken together, substantial evidence indicates that the Rhabdoviridae family possesses desirable characteristics to drive tumor regression within a single cycle, with the capacity to undergo multiple treatment cycles delineated as a current therapeutic limitation.

Here, we hypothesized that the intrinsic serological diversity across the vesiculovirus genus could be exploited to engineer a platform of OVs capable of evading cross-neutralization and enabling repeated dosing, even in the setting of pre-existing humoral immunity. To achieve this, we developed a library of chimeric vesiculoviruses bearing distinct glycoproteins from phylogenetically diverse species, while maintaining the replication-competent VSV backbone. We systematically evaluated this platform for replication fitness, oncolytic potential, neutralization resistance, and capacity to remodel the TME upon repeat intravenous delivery. These studies establish a mechanistically informed, serotype-diverse virotherapy platform that addresses a critical translational barrier in the field and provides a path forward for durable systemic immunovirotherapy in solid tumors.

## Results

### Generation of a Library of Oncolytic Vesiculovirus Vectors

To develop a platform of vesiculovirus vectors suitable for repeat systemic dosing, we first performed a comprehensive phylogenetic analysis of the 22 recognized members of the *vesiculovirus* genus. We assessed sequence diversity at both the amino acid (Fig. 1A) and nucleotide (Sup. Fig. 1A) levels, focusing on the viral glycoproteins, which serve as the principal antigenic targets of neutralizing antibody (nAb) responses. In the clade containing the clinically developed vesicular stomatitis virus (VSV), glycoprotein sequence identity ranged from ∼84% between VSV and Morreton virus (MORV) to ∼37% between VSV and Malpais Spring virus (MSPV) (Sup. Fig. 1B). Based on this phylogenetic analysis, and published evidence of their anti-tumor activity(*23–25*), we selected 7 novel vesiculovirus glycoproteins to engineer a serologically diverse vector library: Morreton (MORV), Carajas (CARV), Radi (RADV), Perinet (PERV), Malpais Spring (MSPV), Isfahan (ISFV), and Jurona (JURV). To maintain compatibility with existing clinical VSV platforms, we generated chimeric viruses by synthetically replacing the VSV Indiana strain glycoprotein open reading frame with one of these alternative vesiculovirus glycoprotein sequences (Fig. 1B), yielding a library of 8 vectors, including the parental VSV, for further characterization.

**Figure 1:**
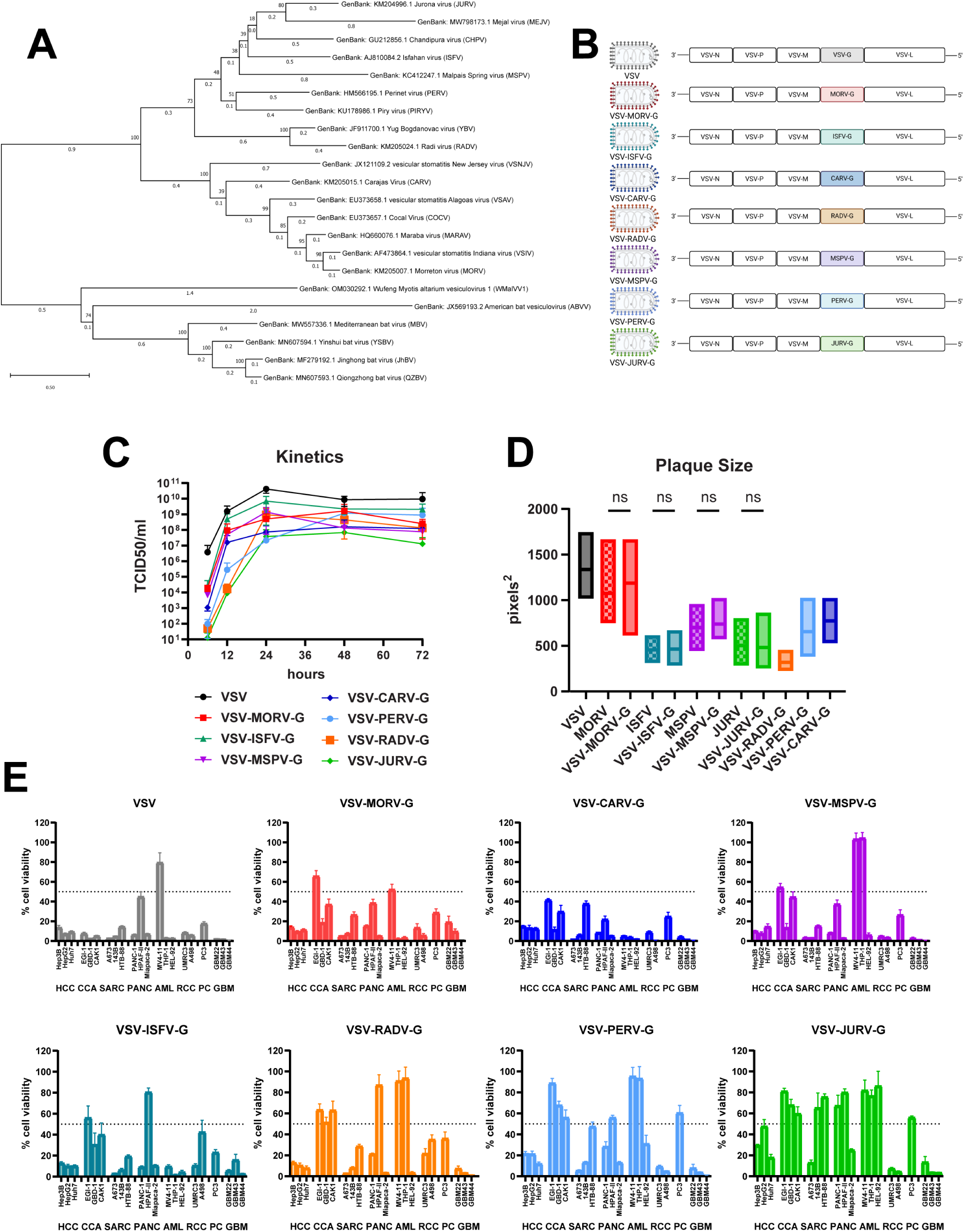
Generation of a Library of Oncolytic Vesiculovirus Vectors. (A) Phylogenetic tree of *vesiculovirus* genus using glycoprotein amino acid sequences to identify evolutionary branching based on viral proteins associated with neutralizing antibodies. Muscle alignment to bootstrap maximum likelihood tree performed in MEGA11 software. (B) Schematic of chimeric vesiculovirus library generation by replacing VSV Indiana glycoprotein with novel vesiculovirus glycoprotein sequence. Chimeric viruses maintain the VSV Indiana nucleocapsid (N), phosphoprotein (P), matrix protein (M), polymerase protein (L) and intergenic regions. (C) Replication kinetics of vesiculovirus library on BHK-21 producer cell line. Following 1-hour infection with MOI 0.1 cell culture supernatants were titered by TCID_50_/ml at respective time points (N=3). (D) Generation of plaque size by wild-type versus chimeric vesiculovirus constructs on Vero-E6 cells. Plaque size was calculated by ViralPlaque macro for ImageJ software (N=3). Significance values determined by ordinary one-way ANOVA with Sidak’s multiple comparisons test. (E) Cell line panel for oncolytic killing by vesiculovirus library. Oncolytic susceptibility was identified as less than 50% cell viability following 72-hour infection as read by MTS reagent (N=9). Outliers were removed from data set using the ROUT method. Tumor cell lines: Hepatocellular carcinoma (HCC), Cholangiocarcinoma (CCA), Sarcoma (SARC), Pancreatic Cancer (PANC), Acute Myeloid Leukemia (AML), Renal Cell Carcinoma (RCC), Prostate Cancer (PC), Glioblastoma (GBM).

All chimeric vesiculoviruses achieved complete lysis of producer cells within 24 hours, generating high titers ranging from 10⁷ to 10⁹ TCID₅₀/mL (Fig. 1C; Sup. Fig. 2). Given the role of the glycoprotein in viral entry, egress, and matrix protein interactions critical for budding, we performed plaque assays to assess replication fitness and cell-to-cell spread. MORV, ISFV, MSPV, and JURV glycoproteins supported plaque formation comparable to their wild-type counterparts (Fig. 1D, Sup. Fig. 3), whereas wild-type RADV did not form plaques, yet the chimeric VSV-RADV-G vector did, highlighting the contribution of the VSV matrix protein in facilitating viral spread (*26–28*). Wild-type CARV and PERV were unavailable for comparison.

We next assessed whether our chimeric vesiculovirus vectors preserved the broad cellular tropism characteristic of VSV, mediated through interactions with the low-density lipoprotein receptor (LDL-R) family (*29*). Across a panel of solid and hematological tumor cell lines, all chimeric vectors induced robust cytopathic effects (Fig. 1E), confirming retained oncolytic activity. However, some cell lines exhibited intrinsic resistance to oncolysis to specific vectors but not others highlighting the inherent advantage of a library of vectors. Therefore, we explored potential mechanisms driving this resistance by measuring type I and III interferon (IFN) expression levels following viral infection. Vesiculoviruses are inherently sensitive to IFN-mediated anti-viral responses, and tumor-intrinsic variability in IFN signaling pathways contributes to differential susceptibility (*30*). Resistant cell lines displayed increased IFN production upon infection compared to susceptible lines (Sup. Fig. 4), aligning with this established mechanism of resistance.

### Vesiculovirus Library Evades Cross Neutralization by VSV Antibodies

To evaluate the vesiculovirus library for cross-neutralization we began with antibodies specific against VSV. As the most clinically advanced candidate, secondary cycles of oncolytic virotherapy are likely to be in the context of pre-existing immunity to VSV. Two monoclonal antibodies, 8G5F11 and 1E9F9, have been characterized to have two distinct epitopes on the VSV glycoprotein (*31*). Neutralization assays with these anti-VSV monoclonal antibodies show distinct dose dependent inhibition of viral cell lysis (Fig. 2A). The IC_50_ value of 8G5F11 is 13 ng/ml and for 1E9F9, is 316 ng/ml. This aligns with previous studies that show that 8G5F11 displays stronger binding than 1E9F9 (*31*). In contrast, each of our synthetic vesiculovirus vectors are uninhibited by anti-VSV monoclonal antibodies demonstrating that these two specific antibody binding epitopes are immunologically distinct between the eight vesiculovirus glycoproteins.

**Figure 2:**
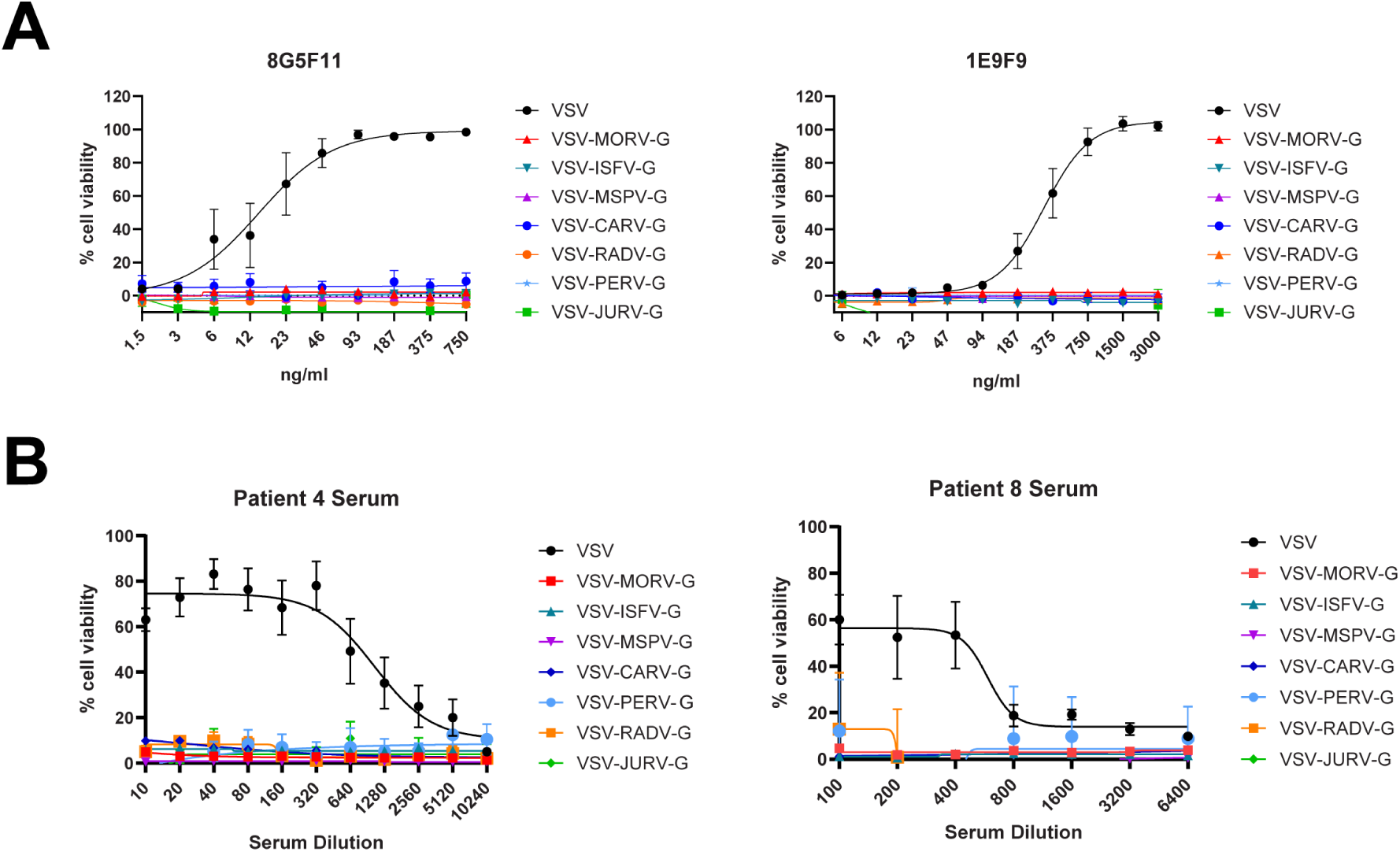
Vesiculovirus Library Evades Cross Neutralization by VSV antibodies. (A) Neutralization assay using commercially available, monoclonal antibodies 8G5F11 and 1E9F9. Virus (MOI 1) and antibody were mixed in equal volumes and incubated for 1-hour at 37°C before plating on BHK-21 cells. Cell viability was read at 72-hours with MTS reagent (N=9). Sigmoidal curve fit with Four-Parameter Logistic Model. (B) Neutralization assay with patient serum from VSV-IFNβ clinical study NCT01628640. Patient serum was first heat inactivated for 30-minutes at 56°C. Virus (500 TCID_50_/units per well) and patient serum dilutions were treated as stated in panel A (N=4).

To further challenge the neutralization evasion capabilities of our library, we performed neutralization assays with sera from hepatocellular carcinoma patients enrolled in a clinical study with VSV-IFNβ oncolytic virotherapy (NCT01628640) (Fig. 2B). Patient sera contain polyclonal antibody populations, which have the capacity to bind to a wide array of epitopes along the VSV glycoprotein in contrast to monoclonal antibodies with singular binding locations. Patient sera containing polyclonal antibody populations against the VSV glycoprotein showed dose dependent neutralization of our VSV vector. However, our vector library completely evaded neutralization by patient sera containing polyclonal anti-VSV antibodies, even at high concentrations. Overall, these data confirm the potential of our vesiculovirus library to evade pre-existing humoral immunity to current VSV clinical candidates and stand as potential clinical candidates for repeat systemic dosing.

### Hypervariability in Epitope Region Models Corroborate Non-Cross Neutralization of Vesiculovirus Library

To elucidate the mechanism of cross-neutralization evasion, we first determined the amino acid sequences of monoclonal antibodies (mAbs) 8G5F11 and 1E9F9 from hybridoma cell lines (courtesy of Dr. Doug Lyles, Wake Forest University). These antibodies were previously used in part to define two major neutralizing epitopes regions on the VSV glycoprotein (Sup. Fig. 5) (*32–36*) For antibody 8G5F11, Minoves et. al, 2025 produced the first high-resolution cryogenic electron microscopy (cryo-EM) structures of full-length VSV-G in complex with the 8G5F11 antibody fragment (Fab) (*37*). No cryo-EM or crystallographic structures currently exist for 1E9F9 in complex with VSV G, leaving its binding mode unresolved.

We then performed deep-learning based de-novo structure predictions using AlphaFold3 (AF3) to model complexes between each antibodies variable heavy (VH) and variable light (VL) chains to the wild-type VSV glycoprotein (*38*). The resulting models recapitulated known epitope mappings: 8G5F11 engaged residues 240–249 within epitope A (Figure 3A), and 1E9F9 targeted residues 358–367 near epitope B (Figure 3B). The 8G5F11 model produced a binding angle of 40° to our trimeric axis, an orientation comparable to the published cryo-EM structures(*37*).

**Figure 3:**
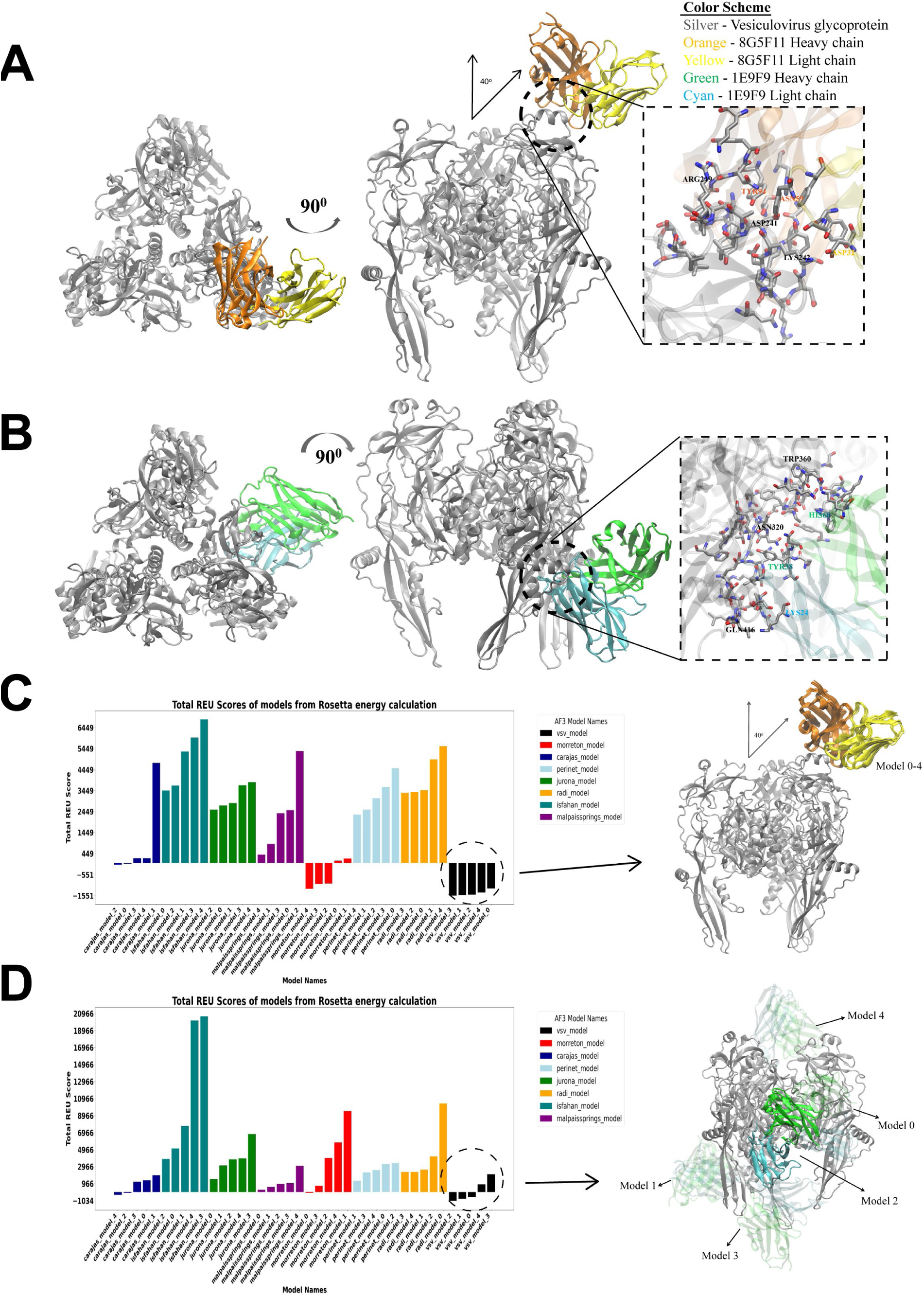
Hypervariability in Epitope Region Models Corroborate Non-Cross Neutralization of Vesiculovirus Library. (A) Top predicted antibody binding model of 8G5F11 to VSV amino acids ADKDLFAAAR. (B) Top predicted antibody binding model of 1E9F9 to VSV amino acids DDWAPYEDVE. (C) Rosetta energy scores of predicted binding interactions between novel vesiculovirus glycoproteins to 8G5F11 mAb. (D) Rosetta energy scores of predicted binding interactions between novel vesiculovirus glycoproteins to 1E9F9 mAb. Lower Rosetta Energy Unit (REU) scores indicate more stable and favorable protein structures. Structural models of the top five docking poses are shown for clarity. All biomolecular structures were visualized and rendered using software Visual Molecular Dynamics (VMD) (*40*).

Subsequently, we generated homo-trimeric models for our novel vesiculovirus glycoproteins using AlphaFold3, which remain uncharacterized by cryo-EM or X-ray crystallography methods, and predicted molecular docking of 8G5F11 and 1E9F9 to each glycoprotein. We ranked the de-novo prediction results using Rosetta software by Rosetta energy scores, where more negative scores represent more favorable protein binding interactions (Figure 3C–D) (*39*). The majority of the novel vesiculovirus glycoproteins produced unfavorable energy scores with either 8G5F11 or 1E9F9, reflecting unstable antibody binding. Among them, MORV glycoprotein yielded favorable scores, a prediction consistent with the close sequence and phylogenetic relationship to VSV. In contrast, although CARV occasionally produced borderline negative scores, these predictions did not translate into observed antibody cross-reactivity.

Importantly, while the cryo-EM structures of 8G5F11 with VSV-G serves as an experimental anchor supporting our docking’s predicted binding orientation, no analogous structural template exists for 1E9F9: the antibody’s binding mode remains computationally inferred only. Together, our AlphaFold-based structural models and Rosetta energy scoring provide a mechanistic rationale for how sequence divergence in key neutralizing epitopes contributes to the evolution of distinct vesiculovirus serotypes that evade cross-neutralization. These computational insights align with our *in vitro* binding assays and support a translational framework for leveraging viral glycoprotein diversity in designing repeat dosing strategies.

### Repeated Intravenous Delivery of Non-Cross Neutralizing Oncolytic Vesiculoviruses Improves Survival in an Immune Competent Murine Cancer Model

We hypothesized that repeat dosing of non-cross neutralized oncolytic vectors would enhance anti-tumor efficacy compared to repeat dosing of the same vector. Therefore, C57BL/6 mice were vaccinated with one of our novel vesiculovirus vectors to generate an anti-viral antibody response (Fig. 4A-B). Following seroconversion and subcutaneous tumor implantation of the B16-OVA-IFNAR^-/-^ cell line, mice received intravenous delivery of a subsequent dose of VSV-OVA, a VSV vector expressing ovalbumin (OVA). We designed the study in this format for multiple reasons: (1) we wanted to generate antibody populations against our novel vesiculovirus glycoproteins, which are not currently commercially available, (2) the B16 model displaying OVA has been shown to respond to VSV-OVA therapy and allows us to differentiate viral antigen directed versus tumor antigen directed T-cell responses (*41*) and (3) the knockout model of the interferon alpha/beta receptor (IFNAR) helps mitigate interferon signaling in the tumor microenvironment and facilitates more robust and durable viral replication. Our primary goal was to assess the antibody populations generated in the mouse. Our secondary objective was to define how pre-existing humoral immunity impacted therapeutic responses to a repeated dose of an oncolytic vesiculovirus vector by intravenous delivery. Effective repeat vector delivery was illustrated through improved survival (Fig. 4C) and reduction in tumor volume (Fig. 4D). In our control group, where we attempted delivery of VSV-OVA in the presence of a pre-existing humoral immunity against the VSV vector, we did not see significant improvement in survival compared to our PBS control group. However, in our treatment groups with non-cross neutralized vectors, there was significant delay in tumor growth achieved through successful delivery of oncolytic virotherapy.

**Figure 4:**
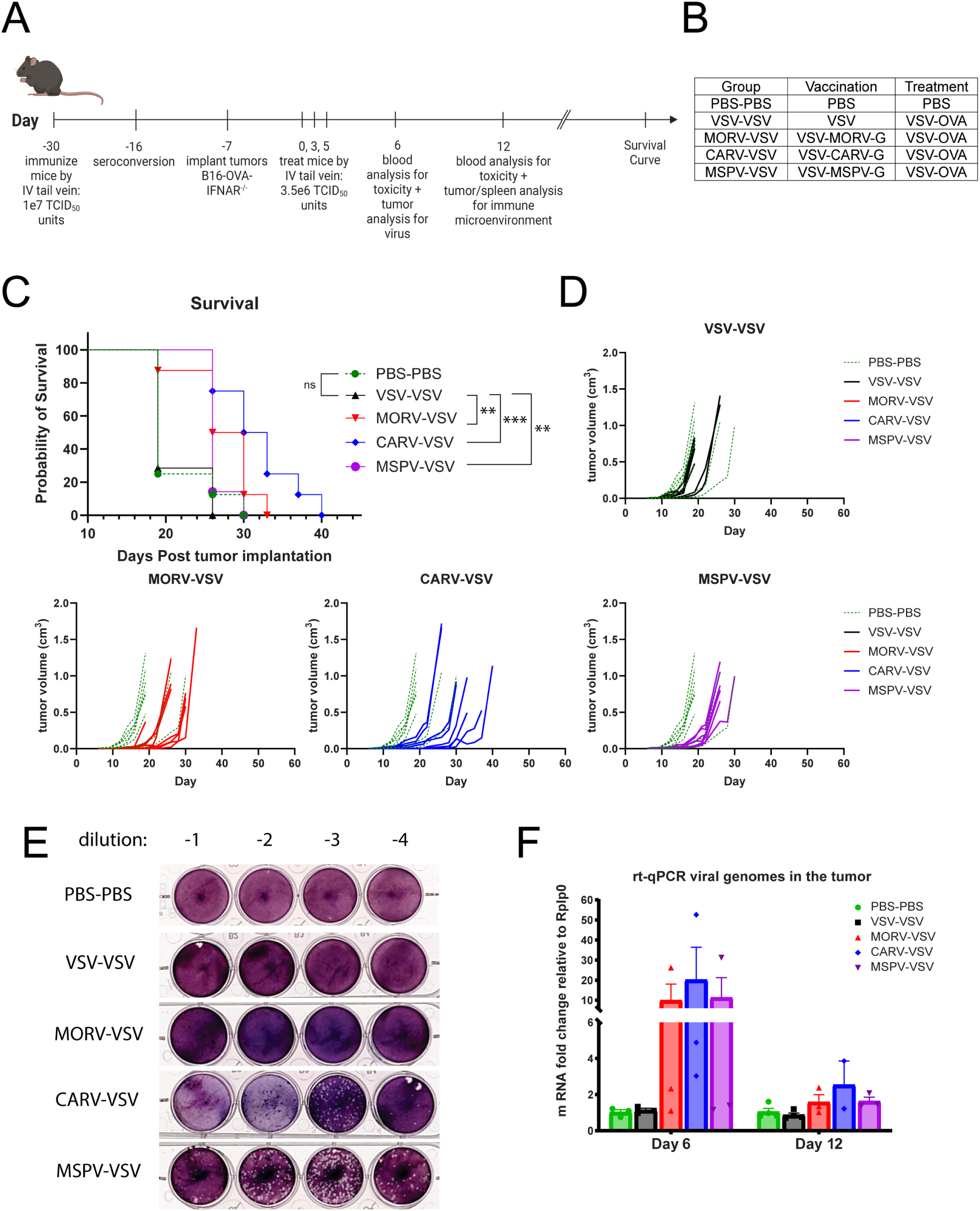
Repeated Intravenous Delivery of Non-Cross Neutralizing Oncolytic Vesiculoviruses Improves Survival in an Immune Competent Murine Cancer Model. (A) Schematic of *in vivo* study design for oncolytic vesiculovirus library. Mice were vaccinated with indicated vesiculovirus vectors and confirmed for seroconversion before tumor implantation. Mice bearing B16-OVA-IFNAR^-/-^ tumors were treated with three intravenous doses of VSV-OVA. One day post treatment cohort mice were examined for toxicity and viral delivery (N=3). Seven days post treatment cohort mice were examined for toxicity, viral delivery, and immune microenvironment changes (N=4). Remaining cohort mice had tumor burden measured until end of study (N=7) (B) Group table for defining vaccination versus treatment virus. (C) Kaplan-Meier survival curve of mice implanted with B16-OVA-IFNAR^-/-^ tumor model. Statistical significance determined by Mantel-Cox test. (D) Tumor volumes of mice implanted with B16-OVA-IFNAR-/- in their respective treatment groups compared to control. (E) Plaque assay of tumors one day post final viral injection. Tumor samples were physically disrupted and lysate underwent 1:10 serial dilutions plated on BHK-21 cells for plaque formation. Plaques indicate live, replication competent virus present in tumor microenvironment. (F) qRT-PCR of viral genomes in the tumor microenvironment one day and seven days post final viral injection.

We confirmed successful viral delivery in the setting of neutralizing antibodies by the recovery of viable viral particles from the tumor in the CARV-VSV and MSPV-VSV groups, observed by the formation of plaques (Fig. 4E). Furthermore, we used quantitative reverse transcription polymerase chain reaction (qRT-PCR) to detect viral RNA in tumors (Fig. 4F). These data show viral delivery in all treatment groups utilizing a novel vesiculovirus vector: MORV-VSV, CARV-VSV, and MSPV-VSV, confirming that it is possible to achieve delivery and therapeutic effect upon second administration of an oncolytic vesiculovirus even in the presence of a potent neutralizing antibody response to an immunologically distinct but related virus.

### Novel Vesiculoviruses Generate Antibody Populations without Cross Reactivity

Next, we tested whether our novel vesiculoviruses produced antibody populations that were cross reactive to our additional vector candidates. We began by confirming seroconversion of our mice 14 days post vaccination (Fig. 5A). The VSV-MSPV-G vector produced a very strong humoral response, with the highest antibody titers. VSV-MORV-G and VSV produced similar antibody titers, but the VSV-CARV-G vector produced notably lower antibody titers. Subsequently, although we have previously shown that anti-VSV sera does not cross react with our vectors (Fig. 2) we tested whether the reverse was true and that VSV would not be neutralized by sera which contained neutralizing antibodies against our novel vector candidates (Fig. 5B). We observed anti-MORV-G antibodies in two of the four mice tested were cross reactive with the VSV vector, albeit, at a lower binding strength as displayed by the high serum dilution necessary to produce a 50% reduction in cell infection. In our VSV-CARV-G and VSV-MSPV-G vaccination groups, antibodies were not cross reactive to VSV.

**Figure 5:**
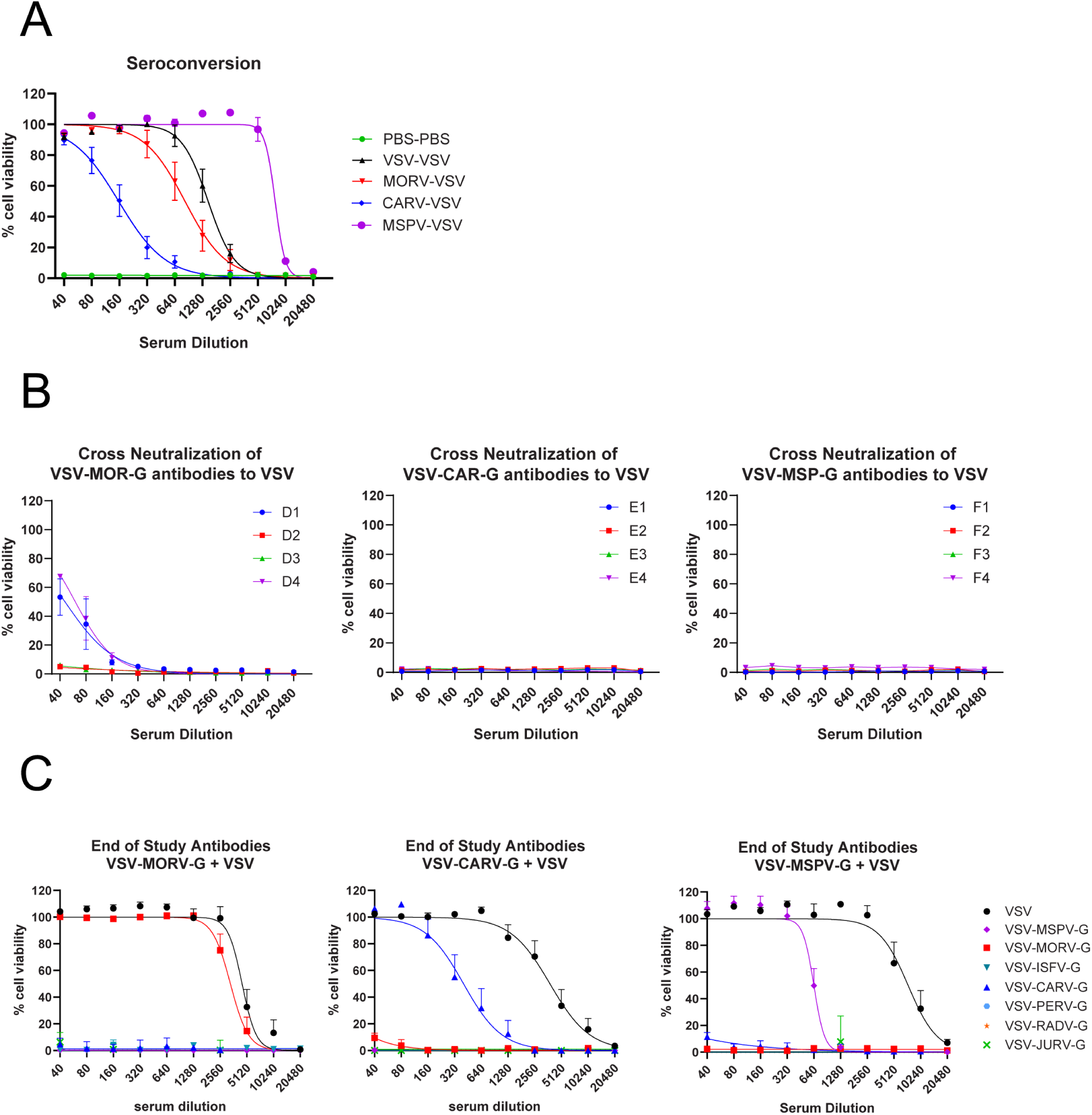
Novel Vesiculoviruses Generate Antibody Populations without Cross Reactivity. (A) Seroconversion of mice vaccinated with oncolytic vesiculovirus. Two weeks post vaccination plasma isolated from cheek bleed was serially diluted and mixed 1:1 with respective vaccination virus (500 TCID50/units). MTS assay readout for cell viability at 72-hours post infection on BHK-21 cells determined antibody titer generated by vaccination (N=4). (B) Cross neutralization of vaccination generated antibodies against VSV vector for treatment post tumor implant. Plasma isolated from cheek bleed indicated in panel A was assessed against VSV vector for potential cross neutralization which would inhibit delivery in tail vein injection of VSV-OVA post tumor implant (N=4). (C) End of study antibody populations were assessed with plasma collected during survival cohort euthanasia. Individual mice per group were assessed against full vesiculovirus library for antibody titers against specific vesiculovirus species glycoproteins (N=4). Sigmoidal curves fit with Four-Parameter Logistic Model.

At the end of the study, we assessed the broad cross-reactivity of antibody populations against the full vesiculovirus vector library (Fig. 5C). In all three treatment groups, we observed distinct antibody populations against the two viral vectors the mice were exposed to, without cross reactivity to any vesiculovirus library member the mouse was naïve to. In a second murine model (CT26-OVA), we assayed the neutralizing antibody responses in BALB/c mice (Sup. Fig 6), which typically have more robust antibody responses compared to C57BL/6 mice. In this model, VSV and VSV-MORV-G antibodies demonstrated greater cross reactivity (Sup. Fig 6E). However, our VSV-CARV-G and VSV-MSPV-G groups maintained distinct antibody populations against the two viral vectors to which the mice were exposed to, without cross reactivity to any vesiculovirus library member to which the mouse was naïve. Therefore, our vesiculovirus vector library has the capacity to generate unique antibody populations, with MORV and VSV exhibiting the only indication for cross-reactivity. This data progresses the evidence for the use of serologically unique vesiculovirus vectors for repeat dosing strategies.

### Repeat Dosing with Heterologous Oncolytic Vesiculovirus Vectors Promotes Tumor Microenvironment Immunomodulation

Clinical evidence has demonstrated that the primary driver of the anti-tumor response to OVs is not direct oncolysis but rather the highly inflammatory nature of viral replication (*42*). This process leads to the release of both viral antigens and tumor-associated antigens into the TME, which in turn primes and activates tumor-specific T-cells, driving systemic anti-tumor immunity, a hallmark of durable therapeutic responses. This immunological shift in the tumor microenvironment is critical for a durable response (*43*). The goal of enabling effective repeat dosing of oncolytic virotherapy in the clinical setting is to prolong TME inflammation and stimulation of immune memory against the tumor for durable therapeutic effect.

We observed consistent and significant shifts towards the induction of pro-inflammatory immune cell populations in the TME following treatments with CARV-VSV, MORV-VSV, and MSPV-VSV when compared to the lack of immune activation in the PBS control group and VSV-VSV treatment group inhibited by humoral immunity (Fig. 6A). Successful delivery of oncolytic virotherapy in the MORV-VSV, CARV-VSV, and MSPV-VSV groups increased numbers of CD3^+^tumor infiltrating lymphocytes (TILs), natural killer cells, pro-inflammatory macrophages, dendritic cells, and B-cells into the TME (Fig. 6A). We further defined our CD3^+^ TIL population as CD8^+^ or CD4^+^ and looked for markers of T-cell activation and exhaustion (Fig. 6B). CD44, a cell surface protein representing T-cell activation and memory T-cells, upregulates on T-cells upon CD3 activation. However, overall protein expression on the cell surface remains lower during rapid clonal expansions and effector function, and is higher during memory function (*44*). Cell surface markers PD-1 and CD39 are associated with T-cell exhaustion when exhibiting high levels of protein expression on the cell surface, but have low level surface expression for crucial T-cell regulation during initial activation (*45*). Therefore, we defined our T-cell populations as activated by increased T-cell counts expressing the cell surface markers of CD44, PD-1, and CD39, while also quantifying the level of protein expression on the cells surface to delineate activation apart from exhaustion. This assessment of T-cell activation versus exhaustion showed that TILs within our novel vesiculovirus treatment groups display T-cell activation and expansion by increased cell counts, without overt exhaustion illustrated by low protein expression, signifying a healthy pro-inflammatory and active tumor microenvironment.

**Figure 6:**
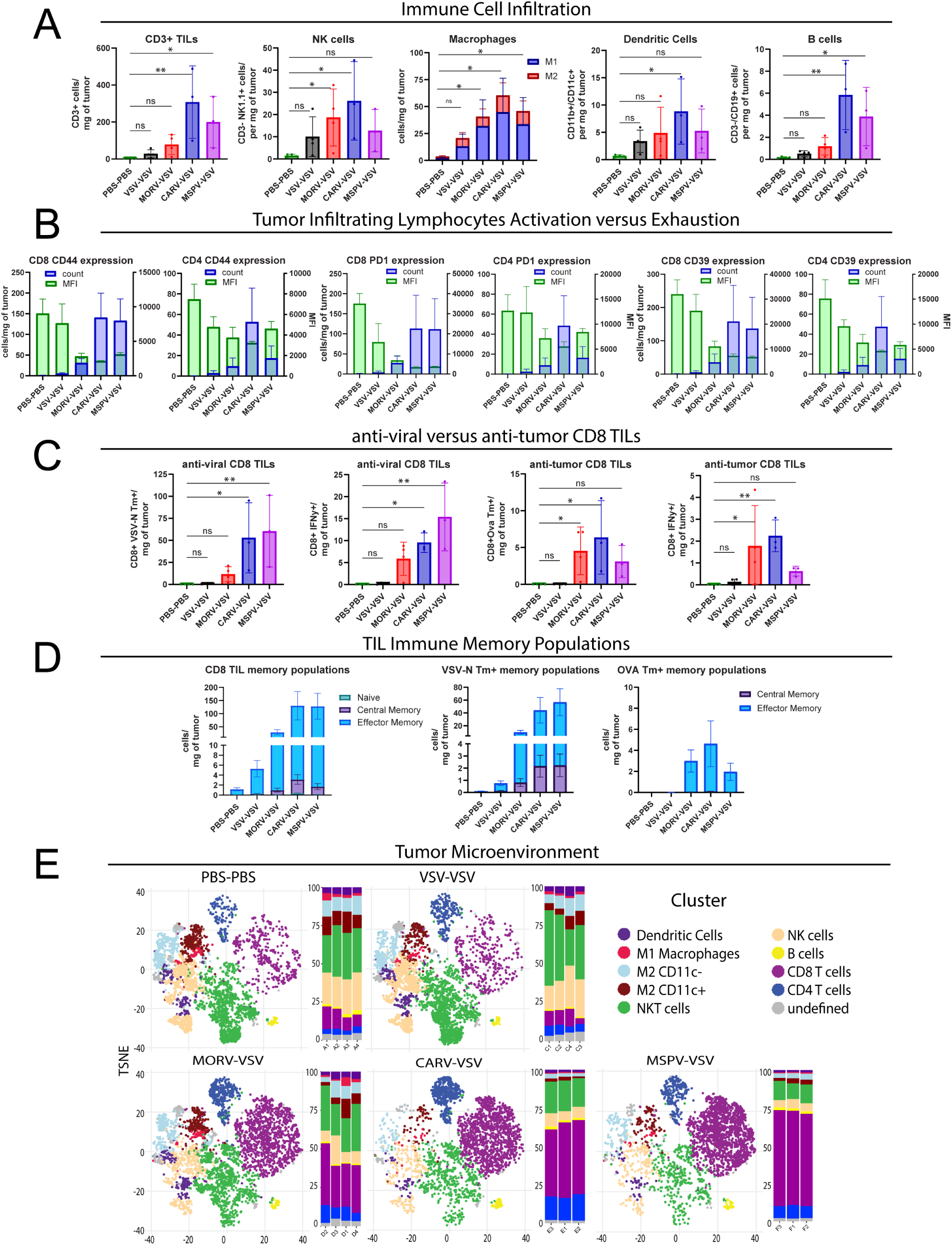
Repeat Dosing with Heterologous Oncolytic Vesiculovirus Vectors Promotes Tumor Microenvironment Immunomodulation. (A) Flow cytometry quantification of immune cell infiltration in the tumor. (B) Flow cytometry quantification of activation and exhaustion markers expressed on CD8^+^ and CD4^+^ T-cell populations. Cell count per milligram of tumor on left Y-axis and median fluorescent intensity (MFI) quantified for defining level of protein expression on cell surface on right Y-axis. (C) Anti-viral and anti-tumor T-cells quantified in two separate experiments utilizing antigen peptides for OVA_257-2640_ (SIINFEKL) and VSV-N_52-59_ (RGYVYQGL). Surface staining was performed with CD8 specific fluorescently labeled MHC tetramers. Intracellular stains were performed with Golgi plug and peptide stimulation before antibody staining. (D) Immune cell memory populations defined by CD44 and CCR7/CD62L co-expression; effector: CD44+CCR7-CD62L-, central: CD44+CCR7+CD62L+ (E) Tumor microenvironment tNSE plots stratified by experimental conditions next to plots of relative abundance of phenotype groups. 2D visualization of cell grouping shows shift in treatment groups towards CD8 and CD4 T-cell infiltration. Statistical significance determined by Kruskal-Wallis test with Dunn’s correction. P values as indicated: ns P > 0.05; * P ≤ 0.05; ** P ≤ 0.01; *** P ≤ 0.001.

We also defined anti-viral versus anti-tumor specific CD8^+^ T-cells by MHCI tetramer staining (Fig. 6C). We classified anti-viral T-cells as T-cells reactive against the VSV nucleocapsid protein epitope N_52-59_, which is present in each of our vesiculovirus library vectors. The tumor specific T-cells were defined by their reactivity against ovalbumin protein epitope OVA_257-264_, which was expressed in both the tumor as a tumor antigen, and by the VSV-OVA vector used to treat the mice. Treatment with MSPV-VSV, which was highly inflammatory based on the strong induction of a humoral response (Fig. 5A), produced a commensurately strong anti-viral T-cell population, which may have been boosted by the therapeutic dose. Interestingly, the MSPV-VSV combination also generated a significantly reduced anti-tumoral (anti-OVA) T-cell response compared to other groups. These data are consistent with a model in which a very strong anti-viral T-cell response can outcompete the development of the anti-tumor T-cell populations (*46*). The CARV-VSV combination, which was associated with the best survival in Figure 4C, generated both strong anti-viral and anti-tumor T-cell responses, suggesting that a Goldilocks balance might be achieved that depends upon the antigenicity of the viral and tumor antigens. Consistent with this ‘balance’ model, the MORV-VSV combination generated a significantly lower anti-viral T-cell response that was accompanied by a relatively strong anti-tumor response (Fig. 6C), despite lower total TILs compared to MSPV-VSV and CARV-VSV (Fig. 6A). This correlated with improved survival in the animals (Fig. 4C). Taken together, these data demonstrate that greater TIL count does not necessarily translate into greater anti-tumor efficacy and highlights the importance of understanding the specificity of infiltrating tumor T-cell responses.

Finally, Figure 6D shows that the majority of memory cells within the TME were effector memory cells (*47, 48*). Furthermore, the anti-viral populations had abundant central memory T-cell responses, consistent with previous immunization and expansion of pre-existing anti-viral memory T-cell groups. In comparison, there were no central memory cells identified in the anti-tumor T-cell groups, suggesting strongly that T-cells reactive against ovalbumin were generated *de novo* in response to the oncolytic virotherapy rather than expansion of pre-existing anti-tumor memory cells resultant of priming by the presence of the tumor itself.

Overall, these data show that repeat dosing with heterologous vesiculovirus oncolytic virotherapy can reverse the immunosuppressive nature of the TME and survival correlates when an optimal balance exists between the recruitment of anti-viral and anti-tumor T-cell responses. Our assessment of the TME shows a shift in immune cell infiltrates between the PBS and VSV-VSV control groups and treatment groups with heterologous viral vector dosing (Fig. 6E). The influx of T-cell populations supports the hypothesis that MORV, CARV, or MSPV immunity does not inhibit the VSV-OVA targeting of, and replication in, the TME. As a result, we observed favorable changes in TME immune dynamics which facilitated a potent anti-viral and anti-tumor response. This TME data supports the use of repeat dosing in oncolytic virotherapy to educate anti-tumor immune cell populations and reinforce tumor regression over time.

## Discussion

We hypothesized that the *vesiculovirus* genus provides a source of glycoproteins with distinct serotypes capable of supporting repeat intravenous dosing by evading humoral immune responses elicited by prior virotherapy. While concerns about VSV-associated neurotoxicity have historically constrained its therapeutic development, attenuation strategies and tumor-targeting approaches have improved safety profiles, supporting its continued clinical evaluation (*10, 49, 50*). We have shown that other vesiculovirus species, such as MORV and JURV, can be used to expand the oncolytic virotherapy platform with safety and efficacy (*23–25*), Therefore, we sought to further leverage this genus by incorporating under-characterized vesiculovirus glycoproteins into the VSV backbone to generate a library of vectors with translational potential for systemic delivery.

RNA viruses induce profound immunological activation within the tumor microenvironment (TME) due to their rapid replication and potent induction of interferon (IFN) responses (*51, 52*). Vesiculoviruses exploit tumor-intrinsic defects in IFN signaling to promote replication and oncolysis, though patient-specific tumor IFN profiles remain a key consideration for clinical deployment (*53*). In the present study, all vectors replicated to high titers within 24 hours but exhibited heterogeneity in plaque morphology, oncolytic potency across tumor cell lines, and capacity to induce IFN responses in resistant lines (Fig. 1C-E; Sup Fig. 4). Although the VSV matrix protein is classically involved in IFN antagonism, inhibition of host transcriptional mechanisms, and resistance to apoptosis (*54–56*), our data demonstrates that glycoprotein switching substantially modulates viral infectivity, spread, and IFN induction (*57, 58*). Importantly, variable direct cell lysis did not correlate with therapeutic potential, aligning with observations that *in vitro* lysis alone does not predict *in vivo* efficacy (*59, 60*). Given the inherent heterogeneity of tumors and their evolving resistance to monotherapies, a library of vesiculovirus vectors with diverse replication and immunogenic profiles could provide a rational strategy to achieve sustained TME remodeling over time (*61–64*).

To counteract the translational barrier posed by humoral immunity, we systematically evaluated the serological distinction of our vesiculovirus library through neutralization assays using patient sera, *in silico* modeling of antibody-glycoprotein interactions, and *in vivo* models assessing evasion of antibody responses. Neutralization assays confirmed VSV Indiana as serologically distinct from our novel glycoprotein vectors (Fig. 2). The 8G5F11 antibody demonstrated high-affinity binding, requiring lower concentrations for neutralization, and our *in silico* modeling corroborated recently published cryo-EM structural data (*37*). In contrast, 1E9F9 exhibited weaker, lateral binding, consistent with its reduced neutralization potency (Fig. 3A-B). The 1E9F9 *in silico* model binding epitope maintained regional consistency with the literature proposed epitope that competes with LDL-R cellular receptor binding (*65*). Using AlphaFold3 and Rosetta energy scoring, we identified major differences in binding energetics that explain the lack of cross-neutralization across our library (Fig. 3C-D). These results highlight the hypervariability of vesiculovirus glycoprotein epitopes. MORV displayed the strongest potential for cross-neutralization, consistent with phylogenetic proximity to VSV, a finding corroborated *in vivo* (Fig. 5B, Sup. Fig. 6E). While this cross-neutralization limited plaque recovery, viral genome detection confirmed successful tumor delivery, and robust anti-tumor immunity was observed (Figure 4). CARV, though phylogenetically close to MORV, maintained sufficient serological distinction to avoid cross-neutralization, further validating the uniqueness of our library (Figure 5).

To enhance therapeutic impact, we incorporated tumor-associated antigens (TAAs) as transgenes within our vectors. Arming OVs with immune-stimulatory transgenes, including TAAs, has emerged as a strategy to potentiate immune responses beyond oncolysis (*66, 67*). Our data showed that the expression of a TAA by our vectors drives potent anti-tumor immunity, even in the presence of pre-existing anti-viral immunity (Fig. 6C-D). Despite pre-established anti-viral T-cells, sufficient activation of tumor-specific T-cells improved survival in the aggressive B16-OVA-IFNAR^-/-^ model. In parallel, primary exposure to a vesiculovirus-TAA vector elicited potent anti-tumor responses (Sup. Fig. 7). These findings support the translational rationale for repeat administration of TAA-expressing vectors to reinforce immune memory and sustain tumor control (*68, 69*).

While we employed a vaccination strategy rather than longitudinal repeat dosing, this choice reflected practical considerations related to survival endpoints in aggressive tumor models and the need to model anti-viral dominance. While neurotoxicity was not assessed in formal toxicology studies, no hindlimb paralysis was observed and systemic analyses showed transient immune activation consistent with viral infection without hepatotoxicity (Sup. Fig. 8A-B). Inflammatory cytokines normalized post-treatment, except in the VSV-VSV group, underscoring the immune impact of homologous re-exposure (Sup. Fig. 8C). Efficacy studies in a CT26-OVA model showed no survival benefit despite preserved serological distinctness, likely due to the aggressive and anergic immune landscape of this model in VSV virotherapy (*70*). Future studies will focus on translationally relevant repeat-dosing schedules (e.g., 2–4 week intervals) and evaluating vector sequencing and its impact on durability of response. Additionally, we appreciate that the use of the non-tolerized OVA antigen in this model is artificial, and our future studies will focus on the use of integrating alternative TAAs and combining OVs with checkpoint inhibitors to prioritize and maximize translational therapeutic potential.

In summary, these findings support a paradigm shift from single-dose oncolytic virotherapy toward multi-dose immunovirotherapy strategies. Clinical studies with rhabdoviruses, including Maraba (NCT02879760) and VSV (NCT03865212), demonstrate that TAA-armed vectors can remodel the TME and expand tumor-specific T-cells. However, concerns regarding neutralizing antibodies and pre-existing anti-viral T-cells limit single-vector strategies. Our serologically distinct vesiculovirus library offers a translationally feasible approach to enable repeat intravenous dosing, sustained tumor engagement, and iterative immune modulation. This novel platform has the potential to advance the field by providing mechanistic and practical foundations for overcoming humoral immunity to achieve durable anti-tumor responses.

## Materials and Methods

### Phylogenetic Analysis

Amino acid or nucleotide sequences were obtained from the National Center for Biotechnology Information (NCBI) reference sequences. For genera information outside of the Vesiculovirus genus, the International Committee on Taxonomy of Viruses (ICTV) virus taxonomy profile: Rhabdoviridae was referenced(*71*). Sequences were aligned using Muscle Alignment tool(*72*). Aligned sequences underwent maximum likelihood fit with 500 bootstrap replicates. Models with the lowest BIC scores (Bayesian Information Criterion) are considered to describe the substitution pattern the best(*73*). Evolutionary analyses were conducted in MEGA11(*74*). Percent Identify matrix performed by Clustal Omega (*75*).

### Generation of Chimeric Viruses

Plasmid pVSV-XN2 underwent laboratory cloning methods to remove VSV-G gene and replace with respective, codon optimized, vesiculovirus G gene (Genscript, USA). To rescue infectious chimeric vesiculoviruses, BHK-21 cells stably expressing T7 RNA polymerase were transfected with full length vesiculovirus genome plasmid and individual helper plasmids expressing VSV-N, VSV-P, and VSV-L using Lipofectamine 3000 (ThermoFisher, USA). Supernatant from P_0_ rescue underwent plaque purification and amplification on BHK-21 cells. Viruses were sucrose gradient purified and sent for sequence validation at Iowa State University College of Veterinary Medicine’s Next Generation Sequencing Core.

### Viral Kinetics

Chimeric vesiculovirus kinetics was performed by seeding 5e5 BHK-21 cells in 6-well plates overnight. The following day cells were infected with an MOI of 0.1 in 1 mL OptiMEM media for 1 hour at 37°C. Infectious media was removed and replaced with 2 mL of complete DMEM. Each well was utilized for a single time point at which time supernatant was removed and titered by the Spearman–Kärber Method for TCID_50_ calculation.

### Viral Plaque Assays

Plaque assays were performed by incubation of 10-fold dilutions of viral stock in 1 mL OptiMEM media on confluent Vero-E6 cells in 6-well plates for 1 hour at 37°C. Infectious media was removed and replaced with 2% agar overlay. 48 hours post infection agar overlay was removed and plaques were visualized by staining with 0.5% crystal violet solution containing 20% methanol. Plates were imaged using GE Amersham Imager 680 and plaque size was calculated using ImageJ macro ViralPlaque (*76*).

### Cancer Cell Line Panel

Cell lines were cultured in ATCC recommended complete DMEM or RPMI. Complete media includes supplementation with 10% Fetal Bovine Serum and 1% Penicillin-Streptomycin. HCC lines: Hep3B, HepG2, Huh7; CCA lines: EGI-1, GBD-1, CAK-1; SARC lines: A673, 143B, HTB-88; PANC lines: PANC-1, HPAF-II, MiaPaca-2; AML lines: MV4-11, THP-1, HEL-92; Borad laboratory stocks of ATCC acquired lines. RCC lines: UMRC3, A498; PC lines: PC3; generous gifts from John Copland Laboratory, Mayo Clinic Florida. GBM lines: GBM22, GBM43, GBM44; generous gifts from Nhan Tran Laboratory, Mayo Clinic Arizona. Cells were seeded at 1e4 cells/well in a 96-well plate in 50 µL complete medium. The following day viruses were added in 10-fold dilutions (MOI 10-0.01) in 50 µL OptiMEM. At 72 hours post infection, cell viability was read according to Promega CellTiter 96® AQueous One Solution Cell Proliferation Assay (MTS) kit instructions. Experiments were performed with 3 technical replicates per plate and 3 experimental replicates.

### Neutralization Assays

Neutralization assays were performed with monoclonal antibodies 8G5F11 and 1E9F9 (Absolute Antibody, UK), patient serum, or mouse plasma. Patient serum and mouse plasma were heat inactivated for 30 minutes at 56°C. Antibody was pre-incubated with virus (500 TCID50/units) in equal volumes for 1 hour at 37°C. Antibody/virus mix was added to BHK-21 cells, seeded day prior at 1e4 cells/well in 96 well plate. 72 hours post infection cell viability was read by Promega CellTiter 96® AQueous One Solution Cell Proliferation Assay (MTS) kit instructions. Experiments were performed with 3 technical replicates per plate. Monoclonal antibody experiments were performed 3 individual times. Patient serum was assessed from two patients at day 22 and 29. Mouse experiments are representative of 4 mice per group.

### Protein – Protein Interaction between Glycoprotein and Monoclonal Antibodies

Independent *de novo* docking calculations were performed using deep learning based AlphaFold3 (AF3) for eight glycoproteins with mAbs 8G5F11 and 1E9F9 (*38*). Only the heavy and light chains of the antibody were used in the complex structure generation protocol with the homo-trimeric glycoprotein protein. Results from AF3 were ranked based on their individual Rosetta energies computed using Rosetta software using energy function REF2015 (*39*). To identify the interacting amino acid residues between monoclonal antibodies and the glycoproteins in the generated structures, we used VMD (*40*) with a standard CAPRI approach, distance cutoff of 5 Å.

Interaction (Ab_i_, p_j_) = 1, if ∃(r_a_∈Ab_i_, r_p_∈p_j_) such that ||x_ra_−x_r_||≤5Å

Interaction (Ab_i_, p_j_) = 0, otherwise.

where Ab_i_ is the *i-th* mAb, p_j_ is the *j-th* glycoprotein, r_a_ and r_p_ are residues in the antibody and viral protein, respectivel. X_ra_ and X_rp_ are the coordinates of the CA atoms in the residues.

### In Vivo Studies

Female C57Bl/6J (Strain #:000664) and BALB/cJ (Strain #000651) were obtained from The Jackson Laboratory (Bar Harbor, ME). Mice were 6-8 weeks of age upon receipt and maintained in a specific pathogen-free BSL2 biohazard facility. *In vivo* studies were done in collaboration with the laboratory of Richard Vile at Mayo Clinic, Rochester. Generous gifts incorporated into the study include tumor models: B16-OVA-IFNAR^-/-^ and CT26-OVA; and viral vectors: VSV-OVA. All animal studies were conducted in accordance with and approved by the Institutional Animal Care and Use Committee at Mayo Clinic.

C57Bl/6J mice were challenged intravenously with respective viral vector at vaccination doses of 1e7 TCID_50_ units in 50ul PBS. Mice were subcutaneously implanted with 5e5 B16-OVA-IFNAR-/- cells. Once tumors reached approximately 0.2cm in diameter, three treatments were administered intravenously at doses of 5e6 TCID_50_/units on day 0, 5e6 TCID_50_/units on day 3, and 1e6 TCID_50_/units on day 5 for an average of 3.5e6 TCID_50_/units per dose. Mice were monitored and tumor volume was calculated using the following equation: (LengthxWidth2)/2.

The CT26-OVA tumor model was generated by transducing the CT26 cell line (ATCC Cat# CRL-2638) with the lentiviral vector pHR-SIIN-ovalbumin-puromycin and maintaining transduced cells in media containing 10ug/ml puromycin (Sigma P9620). BALB/cJ mice were challenged intravenously with respective viral vector at vaccination doses of 5e5 TCID_50_ units in 50ul PBS. Mice were subcutaneously implanted with 5e5 CT26-OVA cells. Once tumors reached approximately 0.2cm in diameter, three treatments were administered intravenously at doses of 2.5e6 TCID_50_/units per dose on day 0, 3, and 5. Mice were monitored and tumor volume was calculated using the following equation: (LengthxWidth2)/2.

### Viral TME Delivery Assays

Resected tumors were individually weighed before being surgically divided for individual analysis by viral plaque assay, qPCR, and flow cytometry. Tumor sections for plaque assay were disrupted in OptiMEM using a Tissueruptor (Qiagen, USA) for no more than 10 seconds to preserve virus integrity. Tissue homogenate was serially diluted in 10-fold volumes before transferring to confluent BHK-21 cells in 12 well plates. Virus infection occurred for 2 hours at 37°C before removal of infection media and overlay with 3% carboxymethyl cellulose (CMC) solution. After 48 hours plaques were visualized by removing CMC overlay and staining cell monolayer with 0.1% crystal violet solution containing 80% methanol.

Viral qPCR was performed by extracting viral RNA from tumor tissues using the RNeasy Mini Kit (Qiagen, USA) following the manufacturer’s protocol. cDNA synthesis was performed using the iScript cDNA Synthesis Kit (Bio-Rad, USA). SYBR Green-based real-time quantitative PCR (qPCR) was conducted using the Power SYBR Green PCR Master Mix (Thermo Fisher Scientific) on a QuantStudio 7 Flex Real-Time PCR System (Applied Biosystems). Expression of VSV-N was quantified using mouse qPCR Primers (forward: 5’-TGTCTACCAAGGCCTCAAATC-3’; reverse: 5’-CCTGCTTTCCCGATGTTTATTC-3’). Gene expression levels were normalized to Rplp0 using mouse qPCR Primers (forward: 5’-CGCTTGTACCCATTGATGATG-3’; reverse: 5’-TTATAACCCTGAAGTGCTCGAC-3’) and analyzed using the ΔΔCt method.

### Flow Cytometry

Single cell suspensions of spleens and tumors were generated from euthanized mice and immediately processed for flow cytometry studies. Tumors were weighed and digested with DNAse I (Sigma) and Liberase TL (Roche) for 30 min in a 37°C water bath. Spleens and tumors were subjected to ACK red blood cell lysis buffer. Cells were stained for surface markers, washed, and fixed in 4% formaldehyde as previously described(*41*). Samples were analyzed in the Mayo Clinic Flow Cytometry Core (Rochester, MN) using a Cytek Aurora spectral flow cytometer with SpectroFlo (V3.1.0) software for unmixing and Flowjo (V10.1) for data analysis. Tetramers for OVA_257-264_ (SIINFEKL)-APC and VSV-N_52-59_ (RGYVYQGL)-PE were obtained from the NIH Tetramer Core Facility. Tetramers were used at a concentration of 1:100 and incubated at 4°C for 30 min before full surface marker panel staining. Murine antibody panel: CD45.2 (eBioscience, NovaFluor 585), CD3 (BioLegend, AF700), CD4 (BioLegend, PerCP), CD8b.2 (BioLegend, PacBlue), CD11b (BioLegend, PerCP/Cy5.5), CD11c (BioLegend, BV510), CD19 (BD Horizon, RB744), NK1.1 (BioLegend, FITC), F4/80 (BioLegend, APC-Fire810), CD206 (BioLegend, PE-Fire700), PD-1 (BioLegend, BV711), CD39 (BioLegend, PE-Dazzle), CCR7 (BioLegend, PE-Cy7), CD44 (BioLegend, BV785), CD62L (BD Horizon, BV605), Zombie live/dead (BioLegend, NIR), IFNy (BioLegend, APC). For intracellular staining and detection of IFNy, cell suspensions were first cultured in RPMI + IL2 (1:1000) at 37°C + 5% CO2 for no more than 12 hours. Suspensions were then stimulated with viral or tumor peptides and Golgi plug 1:1000 (BD) as denoted at a peptide concentration of 1ug/ml for 4 hours. Cells were then stained and fixed using BD Cytofix/Cytoperm plus with BD Golgi Plug kit (BD Biosciences 555028). Peptides were obtained from the Mayo Clinic Proteomics Core Facility (Rochester, MN).

### Statistical Analysis

Statistical analyses were performed using GraphPad Prism 10.3.1 software. Where applicable curves were assessed with Sigmoidal curves with Four-Parameter Logistic Models. *In vitro* data was assessed with One-way ANOVAs with Sidak’s multiple comparison posttest. *In vivo* data was assessed with Kruskal-Wallis test with Dunn’s correction and survival curves were assessed by Mantel-Cox test. P-values have been set at p=<0.05 and ns=>0.05. Error bars denote the SEM of samples. Figures were generated using GraphPad Prism 10.3.1 software, BioRender, and Adobe Illustrator. Cluster plot analysis was performed using R CATALYST package in R.4.3.1. Pre-established exclusion criteria from mouse survival studies included removal of animals with failed tumor engraftment and those which were euthanized due to endpoint criteria outside of tumor sizes reaching 1cm in diameter. Data from spleen samples with poor overall viability were excluded from analysis.

## Acknowledgments

Special thanks to core services provided by the Mayo Clinic which include the Flow Cytometry Core and the Department of Molecular Medicine’s Virus and Gene Therapy Toxicology and Pharmacology Shared Resource. Special thanks to Alexei G. Basnakian, MD, PhD, DSc for analysis of murine liver and kidney toxicology at the University of Arkansas for Medical Sciences Department of Pharmacology and Toxicology at the DNA Damage and Toxicology Core.

## Author contributions

Conceptualization: N.E.; M.B.; A.B

Experiments and Data Acquisition: N.E.; B.K.; S.D.; D.S.; J.V.; J.T.; C.S.; K.F.; K.B.

Methodology: N.E.; B.K.; S.D.; D.S.; J.V.; J.T.; K.F.; K.B.; A.S.; A.B.; R.V.; M.B.

Funding acquisition: M.B.; R.V.

Supervision: D.S.; S.O.; A.J.S.; A.S.; B.N.; A.B.; R.V.; M.B.

Writing – original draft: N.E.

Writing – review & editing: B.K.; D.S.; E.R.; A.C.; C.W.; S.O.; A.J.S.; B.N.; A.B.; R.V.; M.B.

## Data and materials availability

All data are available in the main text or the supplementary materials. Underlying data for computation modeling can be made available upon reasonable request to the corresponding author.

## References

1. A. Melcher, Oncolytic Virotherapy: Single Cycle Cures or Repeat Treatments? (Repeat Dosing Is Crucial!). Mol. Ther. 26, 1875–1876 (2018).

2. S. J. Russell, For the Success of Oncolytic Viruses: Single Cycle Cures or Repeat Treatments? (One Cycle Should Be Enough). Mol. Ther. 26, 1876 (2018).

3. A. Melcher, K. Harrington, R. Vile, Oncolytic virotherapy as immunotherapy. Science 374, 1325–1326 (2021).

4. B. L. Kendall, R. G. Vile, Oncolytic immunovirotherapy: finding the tumor antigen needle in the antiviral haystack. Immunotherapy (2025) (available at https://www.tandfonline.com/doi/abs/10.1080/1750743X.2025.2513853).

5. A. Naing, D. Khalil, O. Rosen, D. R. Camidge, T. Lillie, R.-R. Ji, A. Stacey, M. Thomas, L. Rosen, First-in-human clinical outcomes with NG-350A, an anti-CD40 expressing tumor-selective vector designed to remodel immunosuppressive tumor microenvironments. J. Immunother. Cancer 12, e010016 (2024).

6. D. Mahalingam, C. Fountzilas, J. Moseley, N. Noronha, H. Tran, R. Chakrabarty, G. Selvaggi, M. Coffey, B. Thompson, J. Sarantopoulos, A phase II study of REOLYSIN® (pelareorep) in combination with carboplatin and paclitaxel for patients with advanced malignant melanoma. Cancer Chemother. Pharmacol. 79, 697–703 (2017).

7. J. Lutzky, R. J. Sullivan, J. V. Cohen, Y. Ren, A. Li, R. Haq, Phase 1b study of intravenous coxsackievirus A21 (V937) and ipilimumab for patients with metastatic uveal melanoma. J. Cancer Res. Clin. Oncol. 149, 6059–6066 (2023).

8. A. L. Pecora, N. Rizvi, G. I. Cohen, N. J. Meropol, D. Sterman, J. L. Marshall, S. Goldberg, P. Gross, J. D. O’Neil, W. S. Groene, M. S. Roberts, H. Rabin, M. K. Bamat, R. M. Lorence, Phase I Trial of Intravenous Administration of PV701, an Oncolytic Virus, in Patients With Advanced Solid Cancers. J. Clin. Oncol. 20, 2251–2266 (2002).

9. N. Jayawardena, Poirier, John T, Burga, Laura N, M. and Bostina, Virus–Receptor Interactions and Virus Neutralization: Insights for Oncolytic Virus Development. Oncolytic Virotherapy 9, 1– 15 (2020).

10. J. Cook, K.-W. Peng, T. E. Witzig, S. M. Broski, J. C. Villasboas, J. Paludo, M. Patnaik, V. Rajkumar, A. Dispenzieri, N. Leung, F. Buadi, N. Bennani, S. M. Ansell, L. Zhang, N. Packiriswamy, B. Balakrishnan, B. Brunton, M. Giers, B. Ginos, A. C. Dueck, S. Geyer, M. A. Gertz, R. Warsame, R. S. Go, S. R. Hayman, D. Dingli, S. Kumar, L. Bergsagel, J. L. Munoz, W. Gonsalves, T. Kourelis, E. Muchtar, P. Kapoor, R. A. Kyle, Y. Lin, M. Siddiqui, A. Fonder, M. Hobbs, L. Hwa, S. Naik, S. J. Russell, M. Q. Lacy, Clinical activity of single-dose systemic oncolytic VSV virotherapy in patients with relapsed refractory T-cell lymphoma. Blood Adv. 6, 3268–3279 (2022).

11. J. Lutzky, T. U. Marron, S. F. Powell, D. H. Johnson, M. Patel, A. B. El-Khoueiry, J. Sarantopoulos, S. Dadi-Mehmetaj, L. Russell, S. J. Russell, K. W. Peng, S. Kaesshaefer, G. Gullo, A. S. Bexon, M. Sznol, Optimization of Voyager V1 (VV1) oncolytic virus systemic delivery in combination with cemiplimab and ipilimumab in patients with melanoma and non– small cell lung cancer (NSCLC). J. Clin. Oncol. 40, TPS9595–TPS9595 (2022).

12. K. E. R. Smith, K.-W. Peng, J. S. Pulido, A. J. Weisbrod, C. A. Strand, J. B. Allred, A. N. Newsom, L. Zhang, N. Packiriswamy, T. Kottke, J. M. Tonne, M. Moore, H. N. Montane, L. A. Kottschade, R. R. McWilliams, A. Z. Dudek, Y. Yan, A. Dimou, S. N. Markovic, M. J. Federspiel, R. G. Vile, R. S. Dronca, M. S. Block, A phase I oncolytic virus trial with vesicular stomatitis virus expressing human interferon beta and tyrosinase related protein 1 administered intratumorally and intravenously in uveal melanoma: safety, efficacy, and T cell responses. Front. Immunol. 14 (2023), doi:10.3389/fimmu.2023.1279387.

13. F. Tzelepis, H. K. Birdi, A. Jirovec, S. Boscardin, C. Tanese de Souza, M. Hooshyar, A. Chen, K. Sutherland, R. J. Parks, J. Werier, J.-S. Diallo, Oncolytic Rhabdovirus Vaccine Boosts Chimeric Anti-DEC205 Priming for Effective Cancer Immunotherapy. Mol. Ther. - Oncolytics 19, 240–252 (2020).

14. D. J. Jonker, S. J. Hotte, A. R. Abdul Razak, D. J. Renouf, B. Lichty, J. C. Bell, J. Powers, C. J. Breitbach, D. F. Stojdl, K. B. Stephenson, J. L. Bramson, J. Hummel, C. G. Lemay, J.-C. Cutz, J. Wells, R. Eady, X. Sun, D. Tu, J. Dancey, Phase I study of oncolytic virus (OV) MG1 maraba/MAGE-A3 (MG1MA3), with and without transgenic MAGE-A3 adenovirus vaccine (AdMA3) in incurable advanced/metastatic MAGE-A3-expressing solid tumours: CCTG IND.214. J. Clin. Oncol. 35, e14637–e14637 (2017).

15. E. Ilett, T. Kottke, J. Thompson, K. Rajani, S. Zaidi, L. Evgin, M. Coffey, C. Ralph, R. Diaz, H. Pandha, K. Harrington, P. Selby, R. Bram, A. Melcher, R. Vile, Prime-boost using separate oncolytic viruses in combination with checkpoint blockade improves anti-tumour therapy. Gene Ther. 24, 21–30 (2017).

16. B. W. Bridle, J. E. Boudreau, B. D. Lichty, J. Brunellière, K. Stephenson, S. Koshy, J. L. Bramson, Y. Wan, Vesicular Stomatitis Virus as a Novel Cancer Vaccine Vector to Prime Antitumor Immunity Amenable to Rapid Boosting With Adenovirus. Mol. Ther. 17, 1814–1821 (2009).

17. B. W. Bridle, K. B. Stephenson, J. E. Boudreau, S. Koshy, N. Kazdhan, E. Pullenayegum, J. Brunellière, J. L. Bramson, B. D. Lichty, Y. Wan, Potentiating Cancer Immunotherapy Using an Oncolytic Virus. Mol. Ther. 18, 1430–1439 (2010).

18. J. M. Ricca, A. Oseledchyk, T. Walther, C. Liu, L. Mangarin, T. Merghoub, J. D. Wolchok, D. Zamarin, Pre-existing Immunity to Oncolytic Virus Potentiates Its Immunotherapeutic Efficacy. Mol. Ther. 26, 1008–1019 (2018).

19. E. S. Lambright, E. H. Kang, S. Force, M. Lanuti, D. Caparrelli, L. R. Kaiser, S. M. Albelda, K. L. Molnar-Kimber, Effect of Preexisting Anti-Herpes Immunity on the Efficacy of Herpes Simplex Viral Therapy in a Murine Intraperitoneal Tumor Model. Mol. Ther. 2, 387–393 (2000).

20. D. Dhar, J. F. Spencer, K. Toth, W. S. M. Wold, Effect of Preexisting Immunity on Oncolytic Adenovirus Vector INGN 007 Antitumor Efficacy in Immunocompetent and Immunosuppressed Syrian Hamsters. J. Virol. (2009), doi:10.1128/jvi.02127-08.

21. F. Galivo, R. M. Diaz, P. Wongthida, J. Thompson, T. Kottke, G. Barber, A. Melcher, R. Vile, Single-cycle viral gene expression, rather than progressive replication and oncolysis, is required for VSV therapy of B16 melanoma. Gene Ther. 17, 158–170 (2010).

22. R. J. Prestwich, E. J. Ilett, F. Errington, R. M. Diaz, L. P. Steele, T. Kottke, J. Thompson, F. Galivo, K. J. Harrington, H. S. Pandha, P. J. Selby, R. G. Vile, A. A. Melcher, Immune-Mediated Antitumor Activity of Reovirus Is Required for Therapy and Is Independent of Direct Viral Oncolysis and Replication. Clin. Cancer Res. 15, 4374–4381 (2009).

23. B. M. Nagalo, Y. Zhou, E. J. Loeuillard, C. Dumbauld, O. Barro, N. M. Elliott, A. T. Baker, M. Arora, J. M. Bogenberger, N. Meurice, J. Petit, P. L. S. Uson, F. Aslam, E. Raupach, M. Gabere, A. Basnakian, C. C. Simoes, M. J. Cannon, S. R. Post, K. Buetow, J. C. Chamcheu, M. T. Barrett, D. G. Duda, B. Jacobs, R. Vile, M. A. Barry, L. R. Roberts, S. Ilyas, M. J. Borad, Characterization of Morreton virus as an oncolytic virotherapy platform for liver cancers. Hepatol. Baltim. Md 77, 1943–1957 (2023).

24. C. R. Watters, O. Barro, N. M. Elliott, Y. Zhou, M. Gabere, E. Raupach, A. T. Baker, M. T. Barrett, K. H. Buetow, B. Jacobs, M. Seetharam, M. J. Borad, B. M. Nagalo, Multi-modal efficacy of a chimeric vesiculovirus expressing the Morreton glycoprotein in sarcoma. Mol. Ther. Oncolytics 29, 4–14 (2023).

25. M. Z. Tesfay, Y. Zhang, K. U. Ferdous, M. A. Taylor, A. Cios, R. S. Shelton, C. C. Simoes, C. R. Watters, O. Barro, N. M. Elliott, B. Mustafa, J. C. Chamcheu, A. L. Graham, C. L. Washam, D. Alkam, A. Gies, S. D. Byrum, E. Giorgakis, S. R. Post, T. Kelly, J. Ying, O. Moaven, C. Y. Chabu, M. E. Fernandez-Zapico, D. G. Duda, L. R. Roberts, R. Govindarajan, M. J. Borad, M. J. Cannon, A. G. Basnakian, B. M. Nagalo, Enhancing immune response and survival in hepatocellular carcinoma with novel oncolytic Jurona virus and immune checkpoint blockade. Mol. Ther. Oncol. 32 (2024), doi:10.1016/j.omton.2024.200913.

26. M. M. Ahmed, O. J. Okesanya, B. M. Ukoaka, A. M. Ibrahim, D. E. Lucero-Prisno, Vesicular Stomatitis Virus: Insights into Pathogenesis, Immune Evasion, and Technological Innovations in Oncolytic and Vaccine Development. Viruses 16, 1933 (2024).

27. B. Dancho, M. O. McKenzie, J. H. Connor, D. S. Lyles, Vesicular Stomatitis Virus Matrix Protein Mutations That Affect Association with Host Membranes and Viral Nucleocapsids. J. Biol. Chem. 284, 4500–4509 (2009).

28. T. K. Soh, S. P. J. Whelan, Tracking the Fate of Genetically Distinct Vesicular Stomatitis Virus Matrix Proteins Highlights the Role for Late Domains in Assembly. J. Virol. 89, 11750– 11760 (2015).

29. D. Finkelshtein, A. Werman, D. Novick, S. Barak, M. Rubinstein, LDL receptor and its family members serve as the cellular receptors for vesicular stomatitis virus. Proc. Natl. Acad. Sci. 110, 7306–7311 (2013).

30. S. A. Felt, M. J. Moerdyk-Schauwecker, V. Z. Grdzelishvili, Induction of apoptosis in pancreatic cancer cells by vesicular stomatitis virus. Virology 474, 163–173 (2015).

31. A. M. Munis, M. Tijani, M. Hassall, G. Mattiuzzo, M. K. Collins, Y. Takeuchi, Characterization of Antibody Interactions with the G Protein of Vesicular Stomatitis Virus Indiana Strain and Other Vesiculovirus G Proteins. J. Virol. (2018), doi:10.1128/jvi.00900-18.

32. L. Lefrancois, D. S. Lyles, The interaction of antibody with the major surface glycoprotein of vesicular stomatitis virus I. Analysis of neutralizing epitopes with monoclonal antibodies. Virology 121, 157–167 (1982).

33. W. A. Volk, R. M. Synder, D. C. Benjamin, R. R. Wagner, Monoclonal antibodies to the glycoprotein of vesicular stomatitis virus: comparative neutralizing activity. J. Virol. (1982), doi:10.1128/jvi.42.1.220-227.1982.

34. L. Lefrancois, D. S. Lyles, Antigenic determinants of vesicular stomatitis virus: analysis with antigenic variants. J. Immunol. 130, 394–398 (1983).

35. S. B. Vandepol, L. Lefrancois, J. J. Holland, Sequences of the major antibody binding epitopes of the Indiana serotype of vesicular stomatitis virus. Virology 148, 312–325 (1986).

36. W. Keil, R. R. Wagner, Epitope mapping by deletion mutants and chimeras of two vesicular stomatitis virus glycoprotein genes expressed by a vaccinia virus vector. Virology 170, 392–407 (1989).

37. M. Minoves, M. Ouldali, L. Belot, S. Roche, L. Zakardas, G. Schoehn, Y. Gaudin, A. Albertini, Structures of vesicular stomatitis virus glycoprotein G alone and in complex with a neutralizing antibody, 2025.01.13.632142 (2025).

38. J. Abramson, J. Adler, J. Dunger, R. Evans, T. Green, A. Pritzel, O. Ronneberger, L. Willmore, A. J. Ballard, J. Bambrick, S. W. Bodenstein, D. A. Evans, C.-C. Hung, M. O’Neill, D. Reiman, K. Tunyasuvunakool, Z. Wu, A. Žemgulytė, E. Arvaniti, C. Beattie, O. Bertolli, A. Bridgland, A. Cherepanov, M. Congreve, A. I. Cowen-Rivers, A. Cowie, M. Figurnov, F. B. Fuchs, H. Gladman, R. Jain, Y. A. Khan, C. M. R. Low, K. Perlin, A. Potapenko, P. Savy, S. Singh, A. Stecula, A. Thillaisundaram, C. Tong, S. Yakneen, E. D. Zhong, M. Zielinski, A. Žídek, V. Bapst, P. Kohli, M. Jaderberg, D. Hassabis, J. M. Jumper, Accurate structure prediction of biomolecular interactions with AlphaFold 3. Nature 630, 493–500 (2024).

39. R. F. Alford, A. Leaver-Fay, J. R. Jeliazkov, M. J. O’Meara, F. P. DiMaio, H. Park, M. V. Shapovalov, P. D. Renfrew, V. K. Mulligan, K. Kappel, J. W. Labonte, M. S. Pacella, R. Bonneau, P. Bradley, R. L. Jr. Dunbrack, R. Das, D. Baker, B. Kuhlman, T. Kortemme, J. J. Gray, The Rosetta All-Atom Energy Function for Macromolecular Modeling and Design. J. Chem. Theory Comput. 13, 3031–3048 (2017).

40. W. Humphrey, A. Dalke, K. Schulten, VMD: Visual molecular dynamics. J. Mol. Graph. 14, 33–38 (1996).

41. R. M. Diaz, F. Galivo, T. Kottke, P. Wongthida, J. Qiao, J. Thompson, M. Valdes, G. Barber, R. G. Vile, Oncolytic Immunovirotherapy for Melanoma Using Vesicular Stomatitis Virus. Cancer Res. 67, 2840–2848 (2007).

42. M. E. Davola, K. L. and Mossman, Oncolytic viruses: how “lytic” must they be for therapeutic efficacy? OncoImmunology 8, e1581528 (2019).

43. Y. Tian, D. Xie, L. Yang, Engineering strategies to enhance oncolytic viruses in cancer immunotherapy. Signal Transduct. Target. Ther. 7, 117 (2022).

44. J. Zhou, P. Nagarkatti, Y. Zhong, M. Nagarkatti, Characterization of T-Cell Memory Phenotype after In Vitro Expansion of Tumor-infiltrating Lymphocytes from Melanoma Patients. Anticancer Res. 31, 4099–4109 (2011).

45. A. Chow, F. Z. Uddin, M. Liu, A. Dobrin, B. Y. Nabet, L. Mangarin, Y. Lavin, H. Rizvi, S. E. Tischfield, A. Quintanal-Villalonga, J. M. Chan, N. Shah, V. Allaj, P. Manoj, M. Mattar, M. Meneses, R. Landau, M. Ward, A. Kulick, C. Kwong, M. Wierzbicki, J. Yavner, J. Egger, S. S. Chavan, A. Farillas, A. Holland, H. Sridhar, M. Ciampricotti, D. Hirschhorn, X. Guan, A. L. Richards, G. Heller, J. Mansilla-Soto, M. Sadelain, C. A. Klebanoff, M. D. Hellmann, T. Sen, E. de Stanchina, J. D. Wolchok, T. Merghoub, C. M. Rudin, The ectonucleotidase CD39 identifies tumor-reactive CD8+ T cells predictive of immune checkpoint blockade efficacy in human lung cancer. Immunity 56, 93–106.e6 (2023).

46. R. Vile, B. Kendall, O. Liseth, T. Sangsuwannukul, N. Elliott, M. C. Yerovi, J. Thompson, J. Swanson, S. Rizk, R. Diaz, J. Tonne, Immunodominant antiviral T cell responses outcompete immuno-subdominant antitumor responses to reduce the efficacy of oncolytic viroimmunotherapy. Res. Sq., rs.3.rs-6131273 (2025).

47. C. A. Klebanoff, L. Gattinoni, N. P. Restifo, CD8+ T-cell memory in tumor immunology and immunotherapy. Immunol. Rev. 211, 214–224 (2006).

48. C. R. Plumlee, J. J. Obar, S. L. Colpitts, E. R. Jellison, W. N. Haining, L. Lefrancois, K. M. Khanna, Early Effector CD8 T Cells Display Plasticity in Populating the Short-Lived Effector and Memory-Precursor Pools Following Bacterial or Viral Infection. Sci. Rep. 5, 12264 (2015).

49. A. A. Abdelmageed, S. Dewhurst, M. C. Ferran, Employing the Oncolytic Vesicular Stomatitis Virus in Cancer Virotherapy: Resistance and Clinical Considerations. Viruses 17, 16 (2025).

50. Y. Hua, C. Wang, F. Li, Y. Han, D. Zuo, Y. Lv, M. Sun, P. Yuan, R. Yuan, F. Zhang, L. Ma, Y. Wang, H. Wu, G. Zhou, Q. Lin, S. Wang, N. Li, Y. Lu, Phase 1, open-label, multicenter, dose escalation safety and tolerability study of oncolytic virus OVV-01 in advanced solid tumors. J. Immunother. Cancer 13, e011517 (2025).

51. G. A. F. Vitiello, W. A. S. Ferreira, V. C. Cordeiro de Lima, T. da S. Medina, Antiviral Responses in Cancer: Boosting Antitumor Immunity Through Activation of Interferon Pathway in the Tumor Microenvironment. Front. Immunol. 12 (2021), doi:10.3389/fimmu.2021.782852.

52. L. Wang, L. S. Chard Dunmall, Z. Cheng, Y. Wang, Remodeling the tumor microenvironment by oncolytic viruses: beyond oncolysis of tumor cells for cancer treatment. J. Immunother. Cancer 10, e004167 (2022).

53. O. V. Matveeva, P. M. Chumakov, Defects in interferon pathways as potential biomarkers of sensitivity to oncolytic viruses. Rev. Med. Virol. 28, e2008 (2018).

54. E. J. Faul, D. S. Lyles, M. J. Schnell, Interferon Response and Viral Evasion by Members of the Family Rhabdoviridae. Viruses 1, 832–851 (2009).

55. N. Redondo, V. Madan, E. Alvarez, L. Carrasco, Impact of Vesicular Stomatitis Virus M Proteins on Different Cellular Functions. PLoS ONE 10, e0131137 (2015).

56. S. A. Kopecky, M. C. Willingham, D. S. Lyles, Matrix Protein and Another Viral Component Contribute to Induction of Apoptosis in Cells Infected with Vesicular Stomatitis Virus. J. Virol. 75, 12169–12181 (2001).

57. P. Georgel, Z. Jiang, S. Kunz, E. Janssen, J. Mols, K. Hoebe, S. Bahram, M. B. A. Oldstone, B. Beutler, Vesicular stomatitis virus glycoprotein G activates a specific antiviral Toll-like receptor 4-dependent pathway. Virology 362, 304–313 (2007).

58. E. Jeetendra, K. Ghosh, D. Odell, J. Li, H. P. Ghosh, M. A. Whitt, The Membrane-Proximal Region of Vesicular Stomatitis Virus Glycoprotein G Ectodomain Is Critical for Fusion and Virus Infectivity. J. Virol. 77, 12807–12818 (2003).

59. P. T. Sobol, J. E. Boudreau, K. Stephenson, Y. Wan, B. D. Lichty, K. L. Mossman, Adaptive Antiviral Immunity Is a Determinant of the Therapeutic Success of Oncolytic Virotherapy. Mol. Ther. 19, 335–344 (2011).

60. J. L. Leddon, C.-Y. Chen, M. A. Currier, P.-Y. Wang, F. A. Jung, N. L. Denton, K. M. Cripe, K. B. Haworth, M. A. Arnold, A. C. Gross, T. D. Eubank, W. F. Goins, J. C. Glorioso, J. B. Cohen, P. Grandi, D. A. Hildeman, T. P. Cripe, Oncolytic HSV virotherapy in murine sarcomas differentially triggers an antitumor T-cell response in the absence of virus permissivity. Mol. Ther. - Oncolytics 1, 14010 (2014).

61. S. T. Workenhe, G. Simmons, J. G. Pol, B. D. Lichty, W. P. Halford, K. L. Mossman, Immunogenic HSV-mediated Oncolysis Shapes the Antitumor Immune Response and Contributes to Therapeutic Efficacy. Mol. Ther. 22, 123–131 (2014).

62. T. E. Ginting, J. Suryatenggara, S. Christian, G. Mathew, Proinflammatory response induced by Newcastle disease virus in tumor and normal cells. Oncolytic Virotherapy 6, 21–30 (2017).

63. D. Zamarin, P. Palese, Oncolytic Newcastle Disease Virus for cancer therapy: old challenges and new directions. Future Microbiol. 7, 347–367 (2012).

64. V. Schirrmacher, A. Griesbach, T. Ahlert, Antitumor effects of Newcastle Disease Virus in vivo: Local versus systemic effects. Int. J. Oncol. 18, 945–952 (2001).

65. J. Nikolic, L. Belot, H. Raux, P. Legrand, Y. Gaudin, A. A. Albertini, Structural basis for the recognition of LDL-receptor family members by VSV glycoprotein. Nat. Commun. 9, 1029 (2018).

66. M. K. Melzer, A. Lopez-Martinez, J. Altomonte, Oncolytic Vesicular Stomatitis Virus as a Viro-Immunotherapy: Defeating Cancer with a “Hammer” and “Anvil.” Biomedicines 5, 8 (2017).

67. D. G. Roy, K. Geoffroy, M. Marguerie, S. T. Khan, N. T. Martin, J. Kmiecik, D. Bobbala, A. S. Aitken, C. T. de Souza, K. B. Stephenson, B. D. Lichty, R. C. Auer, D. F. Stojdl, J. C. Bell, M.-C. Bourgeois-Daigneault, Adjuvant oncolytic virotherapy for personalized anti-cancer vaccination. Nat. Commun. 12, 2626 (2021).

68. C. Groeneveldt, J. van den Ende, N. van Montfoort, Preexisting immunity: Barrier or bridge to effective oncolytic virus therapy? Cytokine Growth Factor Rev. 70, 1–12 (2023).

69. J. Martinez-Quintanilla, I. Seah, M. Chua, K. Shah, Oncolytic viruses: overcoming translational challenges. J. Clin. Invest. 129, 1407–1418.

70. N. M. Durham, K. Mulgrew, K. McGlinchey, N. R. Monks, H. Ji, R. Herbst, J. Suzich, S. A. Hammond, E. J. Kelly, Oncolytic VSV Primes Differential Responses to Immuno-oncology Therapy. Mol. Ther. 25, 1917–1932 (2017).

71. P. J. Walker, J. Freitas-Astúa, N. Bejerman, K. R. Blasdell, R. Breyta, R. G. Dietzgen, A. R. Fooks, H. Kondo, G. Kurath, I. V. Kuzmin, P. L. Ramos-González, M. Shi, D. M. Stone, R. B. Tesh, N. Tordo, N. Vasilakis, A. E. Whitfield, ICTV Report Consortium, ICTV Virus Taxonomy Profile: Rhabdoviridae 2022: This article is part of the ICTV Virus Taxonomy Profiles collection. J. Gen. Virol. 103 (2022), doi:10.1099/jgv.0.001689.

72. R. C. Edgar, MUSCLE: multiple sequence alignment with high accuracy and high throughput. Nucleic Acids Res. 32, 1792–1797 (2004).

73. M. Nei, S. Kumar, Molecular Evolution and Phylogenetics (Oxford University Press, 2000).

74. K. Tamura, G. Stecher, S. Kumar, MEGA11: Molecular Evolutionary Genetics Analysis Version 11. Mol. Biol. Evol. 38, 3022–3027 (2021).

75. F. Sievers, A. Wilm, D. Dineen, T. J. Gibson, K. Karplus, W. Li, R. Lopez, H. McWilliam, M. Remmert, J. Söding, J. D. Thompson, D. G. Higgins, Fast, scalable generation of high-quality protein multiple sequence alignments using Clustal Omega. Mol. Syst. Biol. 7, 539 (2011).

76. M. Cacciabue, A. Currá, M. I. Gismondi, ViralPlaque: a Fiji macro for automated assessment of viral plaque statistics. PeerJ 7, e7729 (2019).

