## Supplemental_Material for "Repeat Systemic Delivery of Cross-Neutralization Resistant Synthetic Vesiculoviruses Immunomodulates the Tumor Microenvironment"

### List of Supplementary Materials

### Materials and Methods

#### IncuCyte Apoptosis Assay

96-well flat bottom plates were seeded with 10,000 BHK-21 cells/well in 50µl complete DMEM and allowed to rest overnight. Addition of infection media comprised of 25µl OptiMEM and 1:10 serial dilutions of viral MOI for vector library. Annexin V red for IncuCyte (Essen Bioscience) was added at manufacturers suggested final concentration of 1:200 to a total well volume of 100µl. Plates were sealed with Breathe-Easy membranes (Research Products International; Mt. Prospect, Illinois) and placed into IncuCyte S3 (Sartorius; Germany). Phase and red fluorescence images were taken of wells every 3 hours for a total time of 72 hrs. Images were analyzed in IncuCyte software for red object count per image and results were exported to Excel. Graphs were generated in GraphPad Prism 9 and represent three separate experiments.

**A**

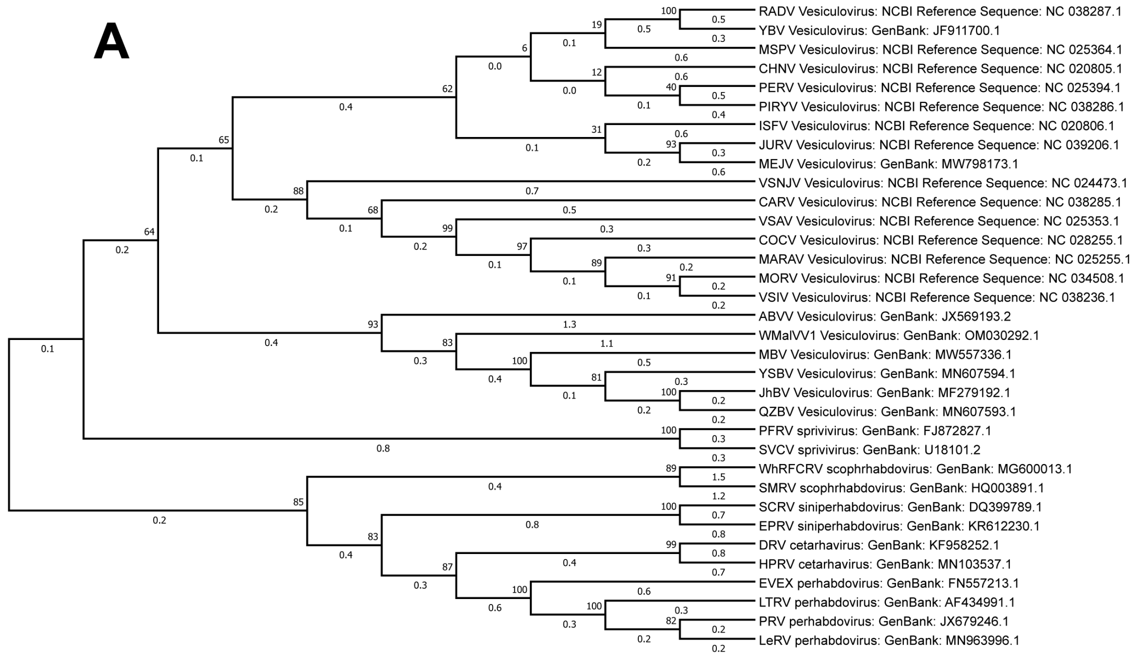


**B**

Percent Identity Matrix - created by Clustal2.1

| | | | | | | | | |
|---|---|---|---|---|---|---|---|---|
| 1: Carajas | 100.00 | 56.73 | 54.99 | 37.52 | 37.96 | 40.00 | 39.07 | 39.34 |
| 2: Morreton | 56.73 | 100.00 | 84.54 | 39.49 | 38.42 | 39.64 | 39.10 | 41.34 |
| 3: VSVindiana | 54.99 | 84.54 | 100.00 | 38.26 | 36.90 | 39.92 | 38.19 | 40.83 |
| 4: Radi | 37.52 | 39.49 | 38.26 | 100.00 | 41.31 | 45.40 | 44.15 | 42.31 |
| 5: MalpaisSpring | 37.96 | 38.42 | 36.90 | 41.31 | 100.00 | 49.81 | 49.90 | 49.13 |
| 6: Perinet | 40.00 | 39.64 | 39.92 | 45.40 | 49.81 | 100.00 | 51.92 | 50.29 |
| 7: Jurona | 39.07 | 39.10 | 38.19 | 44.15 | 49.90 | 51.92 | 100.00 | 54.68 |
| 8: Isfahan | 39.34 | 41.34 | 40.83 | 42.31 | 49.13 | 50.29 | 54.68 | 100.00 |

**Supplemental Figure 1: Nucleotide Phylogenetics and Percent Identity Matrix**

(A) Phylogenetic tree using the glycoprotein nucleotide sequences of the *vesiculovirus* genus, *sprivivirus* genus, *scophrhabdovirus* genus, *siniperhabdovirus* genus, *cetarhavirus* genus, and *perhabdovirus* genus. Allowing evidence of evolutionary changes between viral genera and the glycoprotein, the target of neutralizing antibodies. Muscle alignment to bootstrap maximum likelihood tree performed in MEGA11 software. (B) Percent Identify Matrix performed by Clustal Omega following sequence alignment.

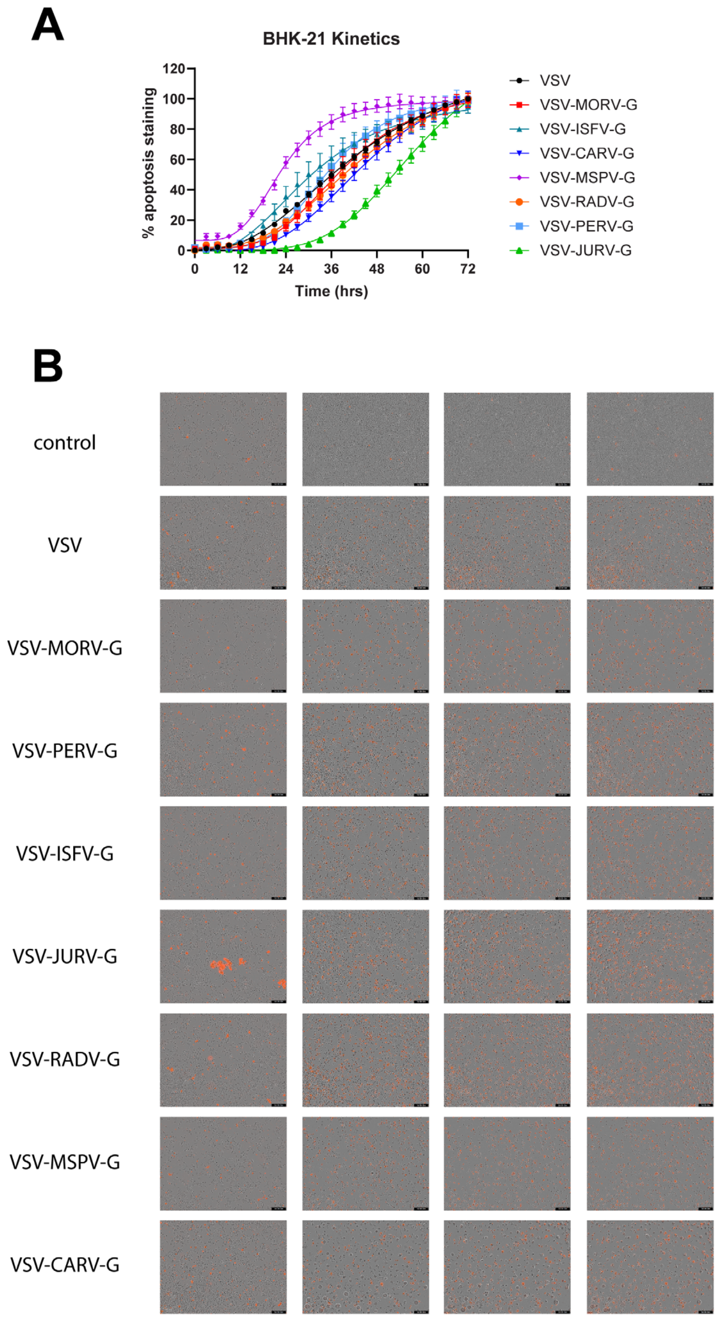


**Supplemental Figure 2: Real-time Annexin V Replication Kinetics**

(A) IncuCyte assay quantitating Annexin V staining of BHK-21 cell death induced by viral lysis at MOI 10 over 72-hours. Images taken every 3 hours. (B) Microscopy images taken by IncuCyte to calculate Annexin V stain at time points 0, 1 day, 2 days, and 3 days.

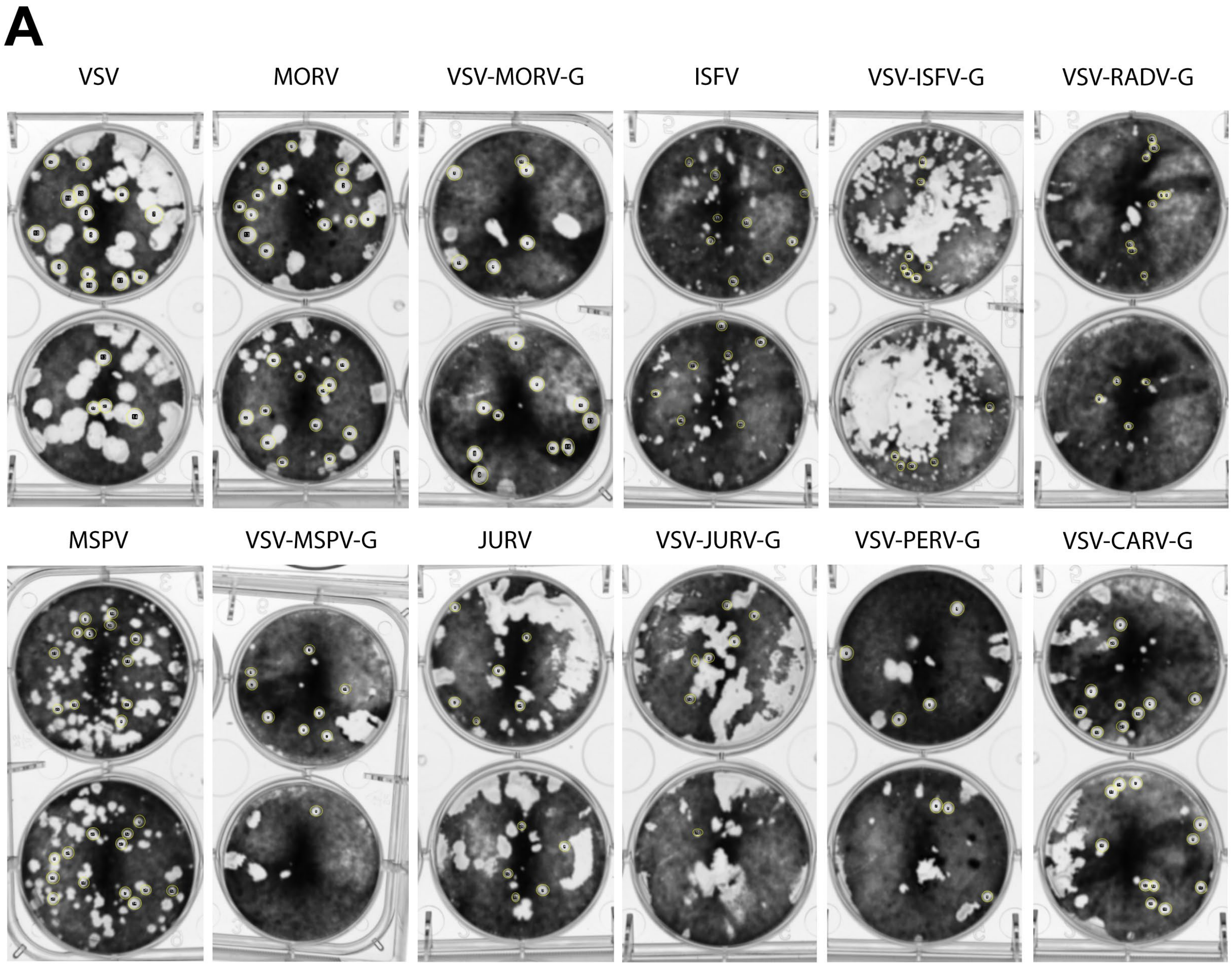


**Supplemental Figure 3: Plaque Assay by ImageJ**

(A) Plaque assay on Vero-E6 cells of wild-type VSV, MORV, ISFV, MSPV, and JURV to compare to plaque size generated by chimeric VSV-MORV-G, VSV-ISFV-G, VSV-MSPV-G, VSV-JURV-G, VSV-RADV-G, VSV-PERV-G, and VSV-CARV-G. Pixel size was determined by ViralPlaque macro installed in ImageJ software (yellow circles).

A

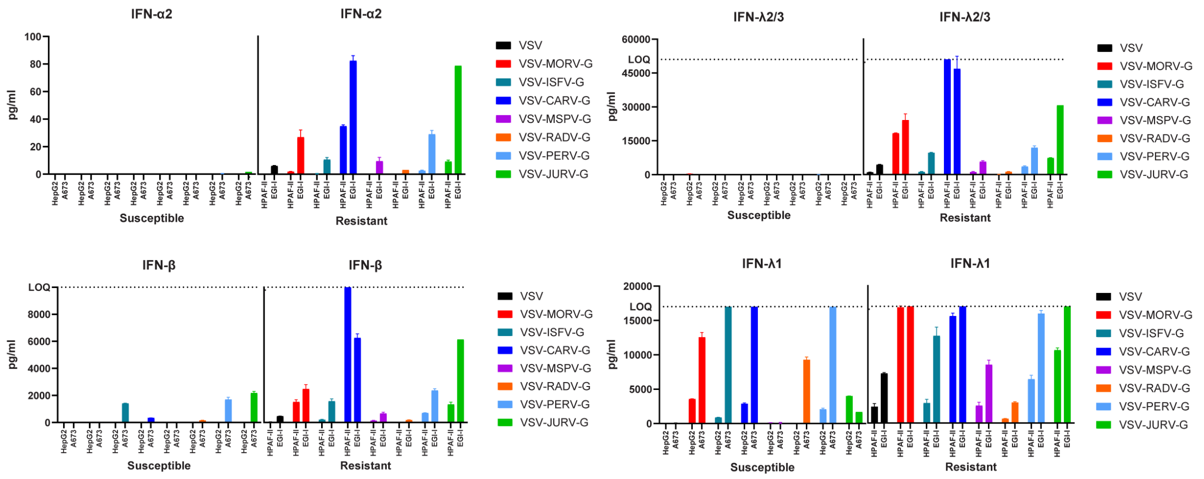


**Supplemental Figure 4: Interferon Signaling of Tumor Cell Lines**

(A) LEGENDplex™ Human Type 1/2/3 Interferon Panel (5-plex) (cat740350) was used to assess cytokine release by previously determined susceptible and resistance cancer cell lines (Figure 1E). Respective cell lines were infected with an MOI 1 for 24 hours before clarifying supernatant and following LegendPlex protocol for analysis of IFN cytokines. Use of LegendPlex data analysis software suite to generate flow cytometry results to pg/ml quantities. Susceptible lines: HCC line HepG2 and SARC line A673; Resistant lines: PANC line HPAF-II and CCA line EGI-1.

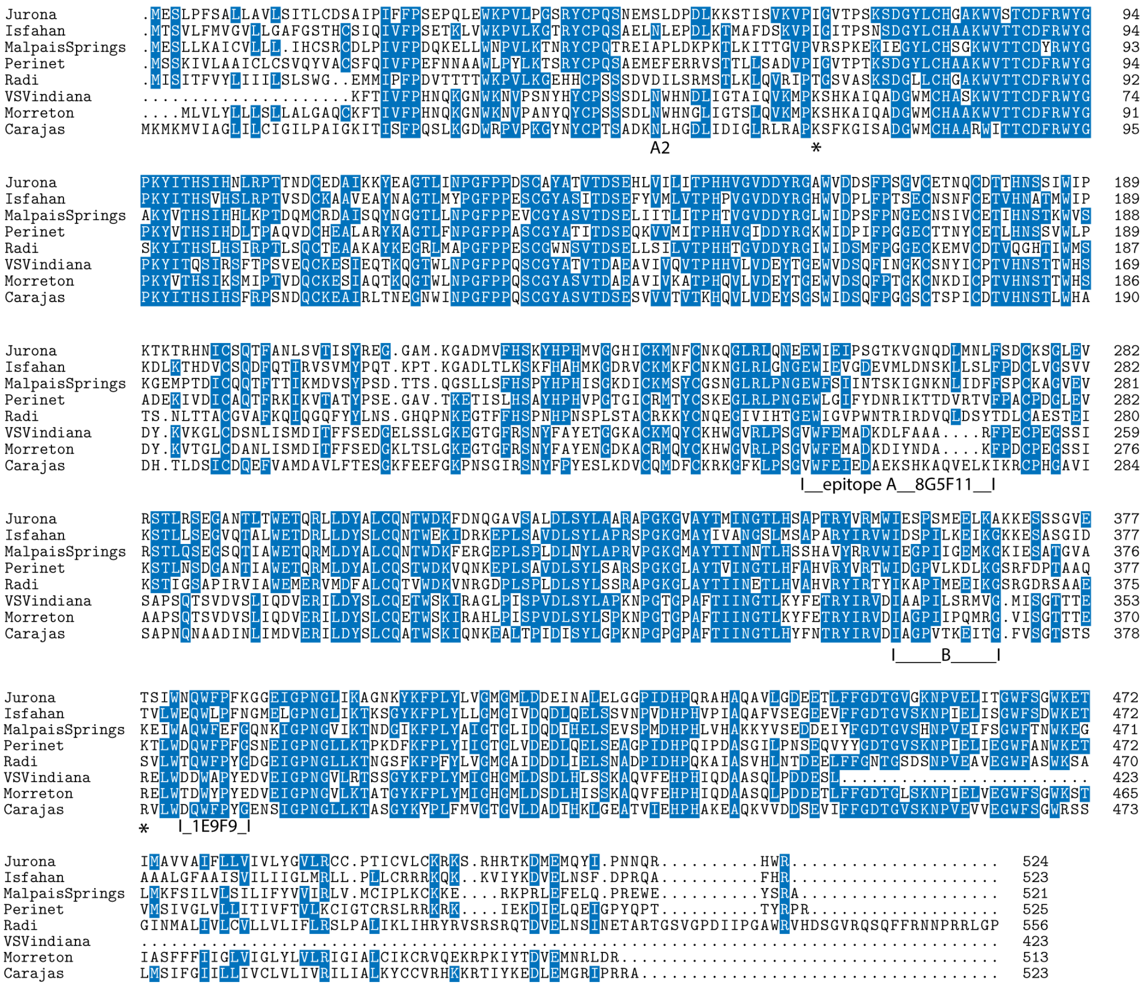


**Supplemental Figure 5: Major Epitope Regions Mapped by 8G5F11 and 1E9F9**

Sequence alignment performed using Clustal W *(75)* with conserved residues highlighted in blue to emphasis regions of conservation versus hypervariability. Two major antibody epitope regions defined as epitope A/A2 and B *(32–36)*. 8G5F11 and 1E9F9 binding defined by *in silico* modeling labeled. * indicates points for ablation of LDL-R binding (*65*). Alignment includes the amino acid residue numbering for reference.

A

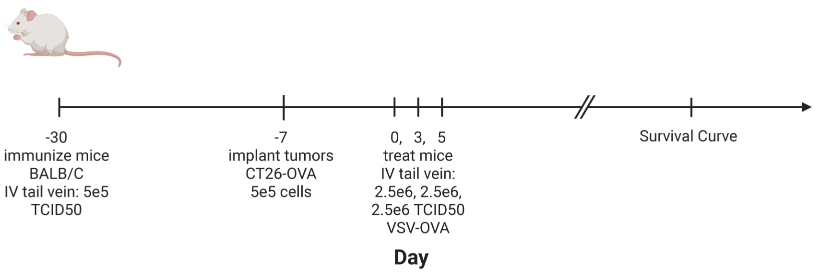


B

| Group | Vaccination | Treatment |
|---|---|---|
| PBS-PBS | PBS | PBS |
| PBS-VSV | PBS | VSV-OVA |
| VSV-VSV | VSV | VSV-OVA |
| MORV-VSV | VSV-MORV-G | VSV-OVA |
| CARV-VSV | VSV-CARV-G | VSV-OVA |
| MSPV-VSV | VSV-MSPV-G | VSV-OVA |

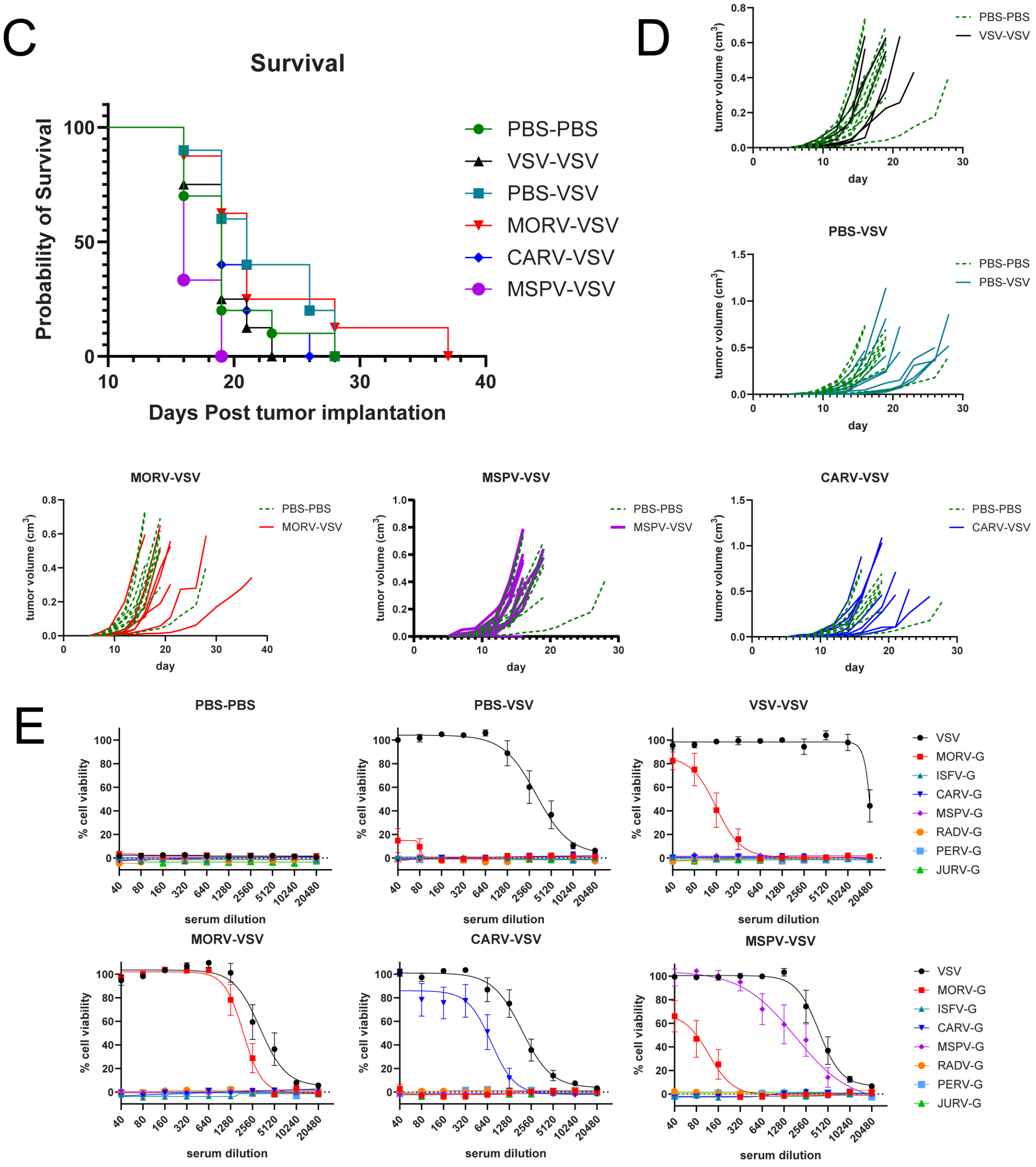

**Supplemental Figure 6: CT26-OVA Model Neutralizing Antibody Response**

(A) *In vivo* study design for secondary murine model. Mice were vaccinated with novel vesiculovirus vector 23 days prior to CT26-OVA tumor implantation. Tumors were allowed to establish for 7 days before three intravenous doses of VSV-OVA. Tumor burden was measured until end of study (N=7). (B) Group table for defining vaccination versus treatment virus. (C) Kaplan-Meier survival curve of mice. Statistical significance determined by Mantel-Cox test. (D) Tumor burdens of mice implanted with CT26-OVA in their respective treatment groups compared to control. (C) End of study antibody populations were assessed with plasma collected during survival cohort euthanasia. Individual mice per group were assessed against full vesiculovirus library for antibody titers against specific vesiculovirus species glycoproteins (N=4). Sigmoidal curves fit with Four-Parameter Logistic Model.

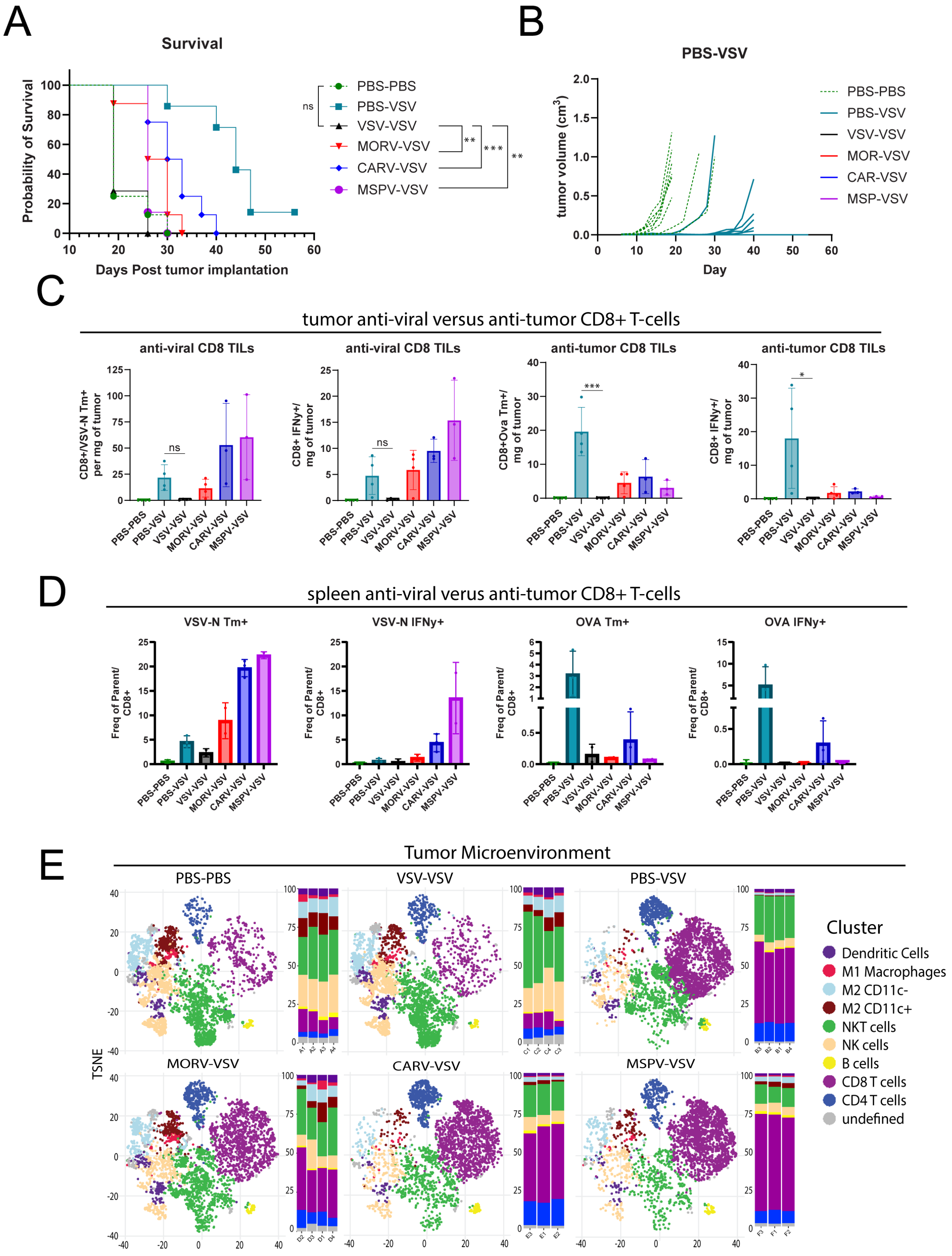
A
Survival
Probability of Survival
Days Post tumor implantation
PBS-PBS
PBS-VSV
VSV-VSV
MORV-VSV
CARV-VSV
MSPV-VSV
ns
**
***
**
B
PBS-VSV
tumor volume (cm3)
Day
PBS-PBS
PBS-VSV
VSV-VSV
MOR-VSV
CAR-VSV
MSP-VSV
C
tumor anti-viral versus anti-tumor CD8+ T-cells
anti-viral CD8 TILs
anti-viral CD8 TILs
anti-tumor CD8 TILs
anti-tumor CD8 TILs
CD8+/VSV-N Tm+ per mg of tumor
CD8+ IFNy+/ mg of tumor
CD8+Ova Tm+/ mg of tumor
CD8+ IFNy+/ mg of tumor
D
spleen anti-viral verus anti-tumor CD8+ T-cells
VSV-N Tm+
VSV-N IFNy+
OVA Tm+
OVA IFNy+
Freq of Parent/ CD8+
E
Tumor Microenvironment
PBS-PBS
VSV-VSV
PBS-VSV
MORV-VSV
CARV-VSV
MSPV-VSV
TSNE
Cluster
Dendritic Cells
M1 Macrophages
M2 CD11c-
M2 CD11c+
NKT cells
NK cells
B cells
CD8 T cells
CD4 T cells
undefined

**Supplemental Figure 7: VSV-OVA group without pre-existing humoral immunity**

(A) Kaplan-Meier survival curve of mice implanted with B16-OVA-IFNAR$^{-/-}$ tumor model. Statistical significance determined by Mantel-Cox test. (B) Tumor burden of mice treated with PBS-VSV treatment group compared to control. (C) Anti-viral and anti-tumor T-cells quantified in two separate experiments utilizing antigen peptides for $OVA_{257\text{-}2640}$ (SIINFEKL) and VSV-$N_{52\text{-}59}$ (RGYVYQGL). External staining was performed with $CD8^+$ specific fluorescently labeled MHCI tetramers. Intracellular stains were performed with Golgi plug and peptide stimulation before antibody staining. (D) splenocytes treated as panel C. (E) Tumor microenvironment tNSE plots stratified by experimental conditions next to plots of relative abundance of phenotype groups. 2D visualization of cell grouping shows shift in treatment groups towards $CD8^+$ and $CD4^+$ T-cell infiltration. Statistical significance determined by Kruskal-Wallis test with Dunn's correction. P values as indicated: ns $P > 0.05$; * $P \leq 0.05$; ** $P \leq 0.01$; *** $P \leq 0.001$.

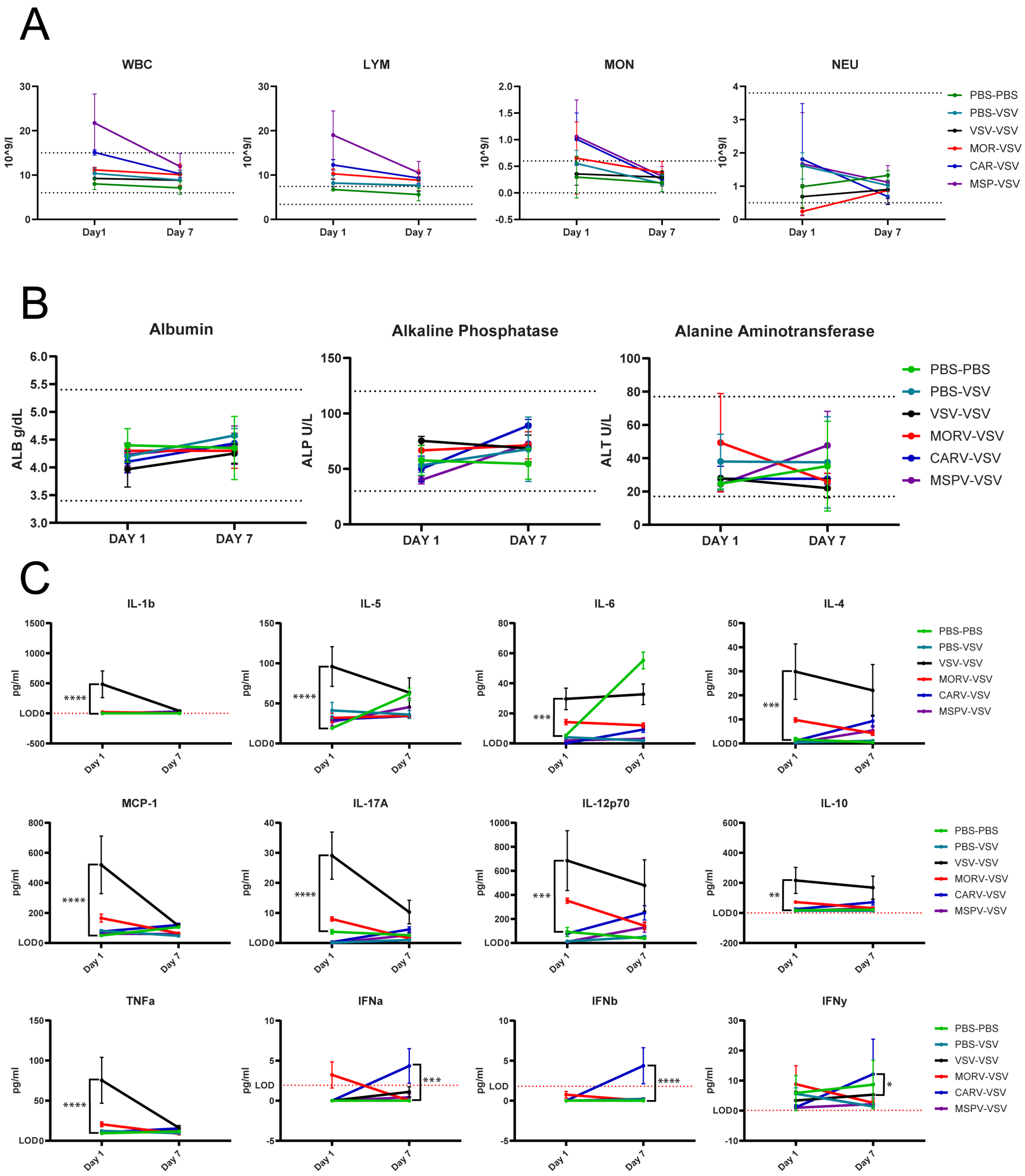
A
WBC
LYM
MON
NEU
10^9/l
Day1
Day 1
Day 7
PBS-PBS
PBS-VSV
VSV-VSV
MOR-VSV
CAR-VSV
MSP-VSV
B
Albumin
Alkaline Phosphatase
Alanine Aminotransferase
ALB g/dL
ALP U/L
ALT U/L
DAY 1
DAY 7
PBS-PBS
PBS-VSV
VSV-VSV
MORV-VSV
CARV-VSV
MSPV-VSV
C
IL-1b
IL-5
IL-6
IL-4
MCP-1
IL-17A
IL-12p70
IL-10
TNFa
IFNa
IFNb
IFNy
pg/ml
LOD
Day 1
Day 7
****
***
**
*
PBS-PBS
PBS-VSV
VSV-VSV
MORV-VSV
CARV-VSV
MSPV-VSV

**Supplemental Figure 8: Blood Toxicity Profile**

(A) Whole blood was immediately aliquoted and assessed by Mayo Clinic's Department of Molecular Medicine Toxicology core. Complete blood count blood analysis was performed on the Abaxis VetScan HM2 hematology analyzer in Rochester, MN. (B) EDTA plasma harvested on respective days was sent to University of Arkansas for Medical Sciences toxicology core for analysis by core director for evidence of liver toxicity. (C) Eve technologies Mouse High Sensitivity 18 plex discovery assay using EDTA plasma harvested on respective days for quantification of inflammatory cytokines expressed in murine blood samples. Eve Technologies Mouse IFN-alpha-beta-2 plex discovery assay using EDTA plasma harvested on respective days for quantification of IFN cytokines in murine blood samples. Statistical significance determined by Mixed-Effects model with Dunnett's multiple comparison test. P values as indicated: ns $P > 0.05$; * $P \leq 0.05$; ** $P \leq 0.01$; *** $P \leq 0.001$; **** $P \leq 0.0001$.

A

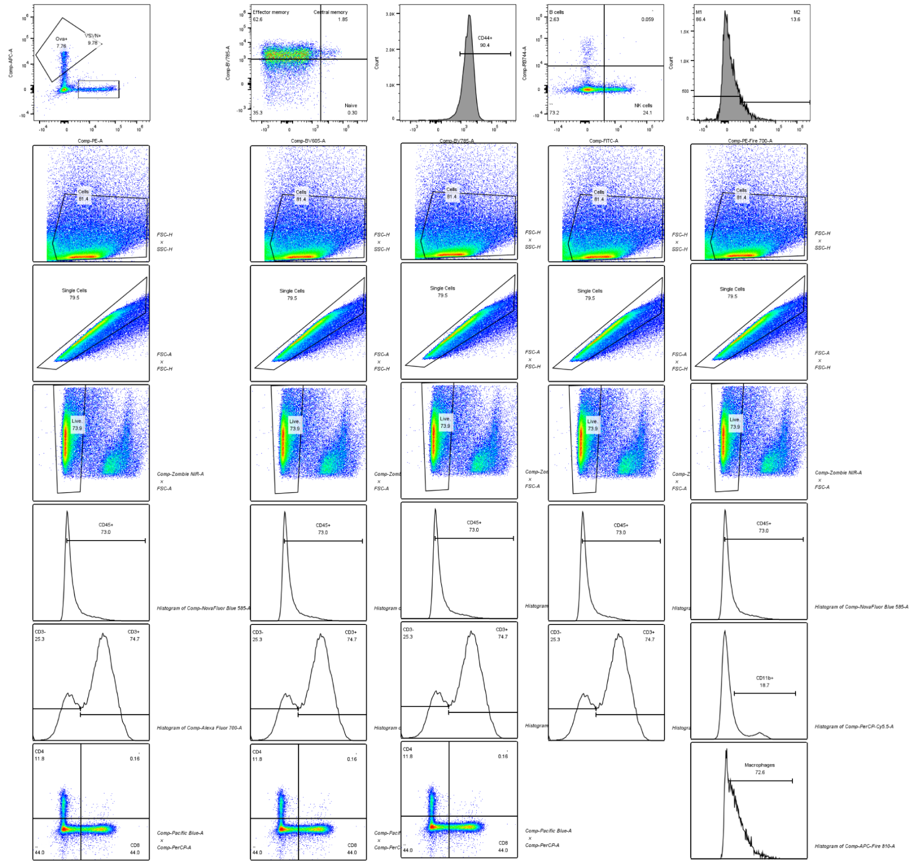

B

| Target | Fluorochrome | Dilution | Company | CAS |
|---|---|---|---|---|
| **Anti-mouse CD8b.2** | PE/Cyanine7 | 1:500 | BioLegend | 140416 |
| **Anti-mouse IFNy** | APC | 1:250 | BioLegend | 505810 |
| **Anti-mouse CD3** | Alexa Fluor 700 | 1:500 | BioLegend | 100216 |
| **Anti-mouse CD45.2** | NovaFluor Blue 585 | 1:350 | eBioscience | M027T02B04 |
| **Anti-mouse CD4** | PerCP | 1:200 | BioLegend | 100538 |
| **Anti-mouse CD8b.2** | Pacific Blue | 1:200 | BioLegend | 140414 |
| **Anti-mouse CD11b** | PerCP/Cyanine5.5 | 1:1000 | BioLegend | 101228 |
| **Anti-mouse CD11c** | Brilliant Violet 510 | 1:350 | BioLegend | 117337 |
| **Anti-mouse CD19** | RB744 | 1:500 | BDHorizon | 570564 |
| **Anti-mouse NK1.1** | FITC | 1:350 | BioLegend | 108705 |
| **Anti-mouse F4/80** | APC-Fire 810 | 1:500 | BioLegend | 123166 |
| **Anti-mouse CD206** | PE-Fire 700 | 1:500 | BioLegend | 141742 |
| **Anti-mouse CD279 (PD-1)** | Brilliant Violet 711 | 1:1000 | BioLegend | 135231 |
| **Anti-mouse CD39** | PE-Dazzle 594 | 1:1000 | BioLegend | 143812 |
| **Anti-mouse CD197 (CCR7)** | PE-Cyanine7 | 1:1000 | BioLegend | 120124 |
| **Anti-mouse CD44** | BV785 | 1:1000 | BioLegend | 103059 |
| **Anti-mouse CD62L** | Brilliant Violet 605 | 1:350 | BDHorizon | 563252 |
| **Live/dead** | Zombie NIR | 1:1500 | BioLegend | 423105 |

**Supplemental Figure 9: Antibodies and Gating Strategy**

(A) Primary gating strategies used to quantify results of tumor microenvironment and spleen immune cell populations. Examples of MHC tetramer stain, effector vs memory populations, activation vs exhaustion markers, and $CD3^+$, B-cell, NK-cell, macrophage, and DC populations. (B) Table of antibodies used in flow cytometry experiments.
